# Enhanced pollen tube performance at high temperature contributes to thermotolerant fruit production in tomato

**DOI:** 10.1101/2024.08.01.606234

**Authors:** Sorel V. Yimga Ouonkap, Meenakshisundaram Palaniappan, Kelsey Pryze, Emma Jong, Mohammad Foteh Ali, Benjamin Styler, Rasha Althiab Almasaud, Alexandria F. Harkey, Robert W. Reid, Ann E. Loraine, Steven E. Smith, Gloria K. Muday, James B. Pease, Ravishankar Palanivelu, Mark A. Johnson

**Affiliations:** Department of Molecular Biology, Cell Biology, and Biochemistry; Brown University; School of Plant Sciences; University of Arizona; Department of Biology, Wake Forest University; Department of Bioinformatics and Genomics; UNC Charlotte; School of Natural Resources and the Environment; University of Arizona; Department of Evolution, Ecology and Organismal Biology; The Ohio State University

## Abstract

Rising temperature extremes during critical reproductive periods threaten the yield of major grain and fruit crops. Flowering plant reproduction depends on development of sufficient numbers of pollen grains and on their ability to generate a cellular extension, the pollen tube, which elongates through the pistil to deliver sperm cells to female gametes for double fertilization. These critical phases of the life cycle are sensitive to temperature and limit productivity under high temperature (HT). Previous studies have investigated the effects of HT on pollen development, but little is known about how HT applied during the pollen tube growth phase affects fertility. Here, we used tomato as a model fruit crop to determine how HT affects the pollen tube growth phase, taking advantage of cultivars noted for fruit production in exceptionally hot growing seasons. We found that exposure to HT solely during the pollen tube growth phase limits fruit biomass and seed set more significantly in thermosensitive cultivars than in thermotolerant cultivars. Importantly, we found that pollen tubes from the thermotolerant Tamaulipas cultivar have enhanced growth *in vivo* and *in vitro* under HT. Analysis of the pollen tube transcriptome’s response to HT allowed us to develop hypotheses for the molecular basis of cellular thermotolerance in the pollen tube and we define two response modes (enhanced induction of stress responses, and higher basal levels of growth pathways repressed by heat stress) associated with reproductive thermotolerance. Importantly, we define key components of the pollen tube stress response identifying enhanced ROS homeostasis and pollen tube callose synthesis and deposition as important components of reproductive thermotolerance in Tamaulipas. Our work identifies the pollen tube growth phase as a viable target to enhance reproductive thermotolerance and delineates key pathways that are altered in crop varieties capable of fruiting under HT conditions.

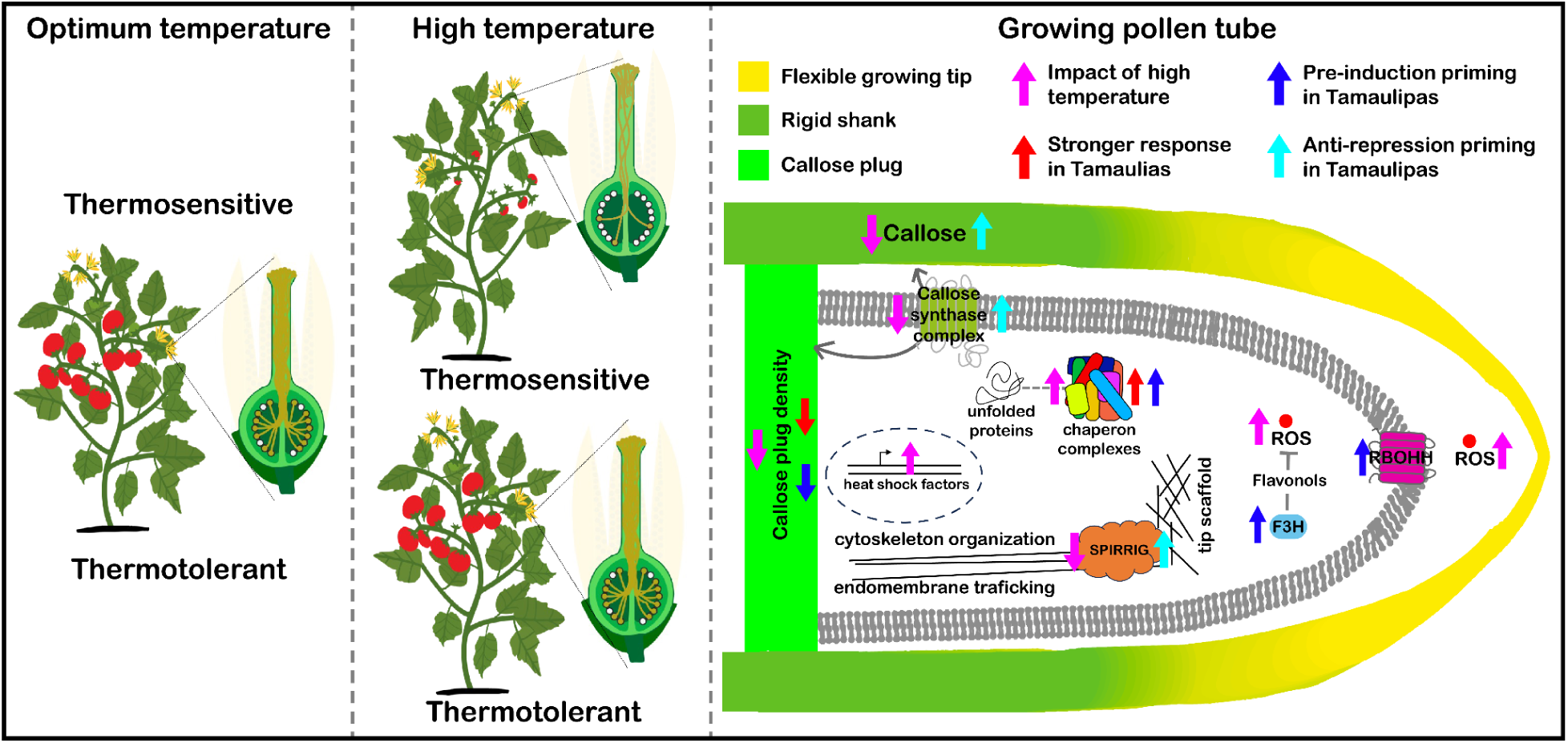

## Introduction

Average daily temperatures have been steadily increasing in recent years and are projected to rise by an estimated average of 3.5° to 6.0°C by the year 2100 ^1^. Agricultural productivity is particularly vulnerable to this change, and rising temperatures are predicted to reduce crop yields. For food crops such as wheat, corn, soybean, rice, and tomato, crop yields are estimated to decrease by 2.5-16% for every additional 1°C of seasonal warming ^2–7^. As a result, research has focused on understanding how plants respond to high temperatures (HT) during vegetative growth ^8–17^ and the development of reproductive cells and structures ^18–39^. The sensitivity of reproductive processes to temperature is of particular interest because of their direct involvement in the production of seed, grain and fruit crops ^2–4,40,41^. To protect our food supply in the face of a warming climate, new methods will be critical to improve crop productivity during HT. A promising strategy is to elucidate the molecular mechanisms that allow thermotolerant plants to produce seeds and fruits even during exceptionally hot growing seasons, and to utilize this knowledge for crop improvement.

Cultivated tomato and its wild relatives offer an excellent model system to understand the genetics of reproductive thermotolerance because natural and artificial selection have led to their adaptation to diverse climates and growth conditions, including extreme temperature variations ^42,43^. Furthermore, field studies have shown a 3°C increase in mean daily temperatures during the reproductive season resulted in a 70% decrease in fruit production in thermosensitive cultivars ^7^. Moreover, genome sequences are available for hundreds of accessions, which will facilitate identification of genetic variants associated with reproductive resilience ^42–45^.

Flowering plant reproduction requires pollen to deliver sperm to female gametes for fertilization, which initiates seed and fruit development ^46^. Pollen grains develop in the anther from hundreds/thousands of diploid mother cells. Each of these cells undergo meiosis to generate tetrads of haploid spores (microsporogenesis, Fig. 1B, left) and each of these spores divides asymmetrically to form a pollen (vegetative) cell with a generative cell in its cytoplasm. In species with tricellular pollen (e.g. Arabidopsis), pollen development is complete when the generative cell divides to form a pair of sperm cells. In bicellular species (e.g. tomato), generative cell division occurs during pollen tube growth ^47,48^. The pollen tube growth phase begins when pollen grains, deposited on the stigma of a compatible pistil, each germinate to form a polarized cellular projection called the pollen tube. Each pollen tube elongates by tip growth ^49^ through specialized transmitting tissue of the style to reach the ovary, which is the site of ovule development. A pollen tube responds to attractants secreted by synergid cells ^50,51^ within each ovule and bursts within one of these synergid cells to release its cargo of two sperm for double fertilization ^52,53^. One sperm cell fuses with the egg to produce the embryo, the other fuses with the central cell to produce endosperm (two major components of seeds); double fertilization also promotes development of the ovary into a fruit. Successful sperm delivery by pollen tubes followed by double fertilization will directly influence the number of seeds per fruit as well as fruit biomass, which are the key components of crop yield ^2,54–56^. Therefore, understanding how HT affects pollen function at the molecular level is a crucial objective.

**Figure 1:**
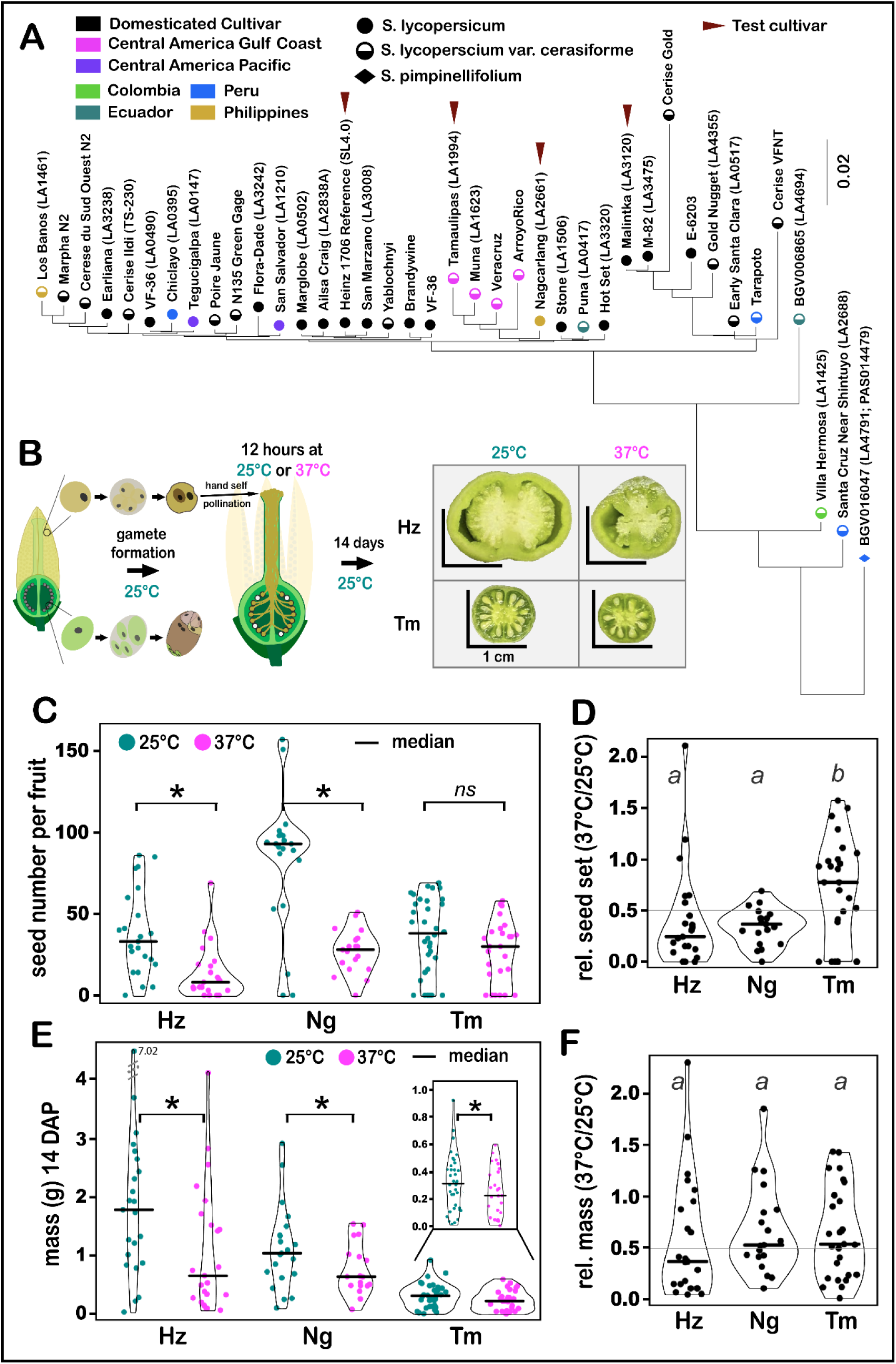
High temperature, applied only during the pollen tube growth phase, decreases seed set and fruit biomass - Tamaulipas is thermotolerant for seed set. **(A)** Phylogenetic tree constructed from whole genome sequences of 39 cultivars of tomato including our four focus cultivars (arrowhead). **(B)** Pollen grains that developed at 25°C were hand pollinated onto emasculated pistils of the same cultivar and incubated for 12 hours at either 25°C or 37°C. Fruit biomass and seed number were measured 14 days after pollination (14 DAP). **(C)** Seed number per fruit. **(D)** Relative seed number (37°C/daily median value at 25°C). **(E)** Fruit biomass (grams) 14 days after hand self pollination. **(F)** Relative fruit biomass (37°C/daily median value at 25°C). Statistical analysis was done using the Mann-Whitney *U* test in C and E (*ns, p >* 0.05; *\**, *p* ≤ 0.05), and using the Kruskal-Wallis test and Dunn’s test for D and F. Similar letters indicate no significant difference (*p* > 0.05) between groups from Dunn’s test. Hz, Heinz; Ng, Nagcarlang; Tm, Tamaulipas.

Research into the effects of HT on plant reproduction has focused on pollen development rather than pollen tube growth. HT has been shown to disrupt anther development ^18,21,22,25,27,31,57^ and limits the number of viable pollen grains produced in the anther ^24^. The impact of HT on meiosis has also been well documented ^58,59^ and HT applied immediately after meiosis also limits pollen production in maize ^57^. Interestingly, some of these negative effects were found to be attenuated in thermotolerant maize cultivars that continued to set fruit at HT ^28,35^. Multiple metabolic and molecular pathways have been identified that respond to HT during pollen development, including the heat shock response and factors that protect against oxidative damage ^26,30,32–37^. An important insight from these studies is that chronic HT limits photosynthetic capacity of the plant, which results in pollen grains with lower accumulation of soluble sugars that are necessary for pollen function ^24,28,29,33^. Notably, this effect was less evident in thermotolerant cultivars ^28^. Despite this progress, cohesive molecular pathways explaining thermosensitivity of plant reproduction remain to be identified ^60^. Therefore, research on how HT affects pollen tube growth in both thermosensitive and thermotolerant cultivars is needed to complement the work being done on pollen development.

Recently, it has been shown that a tomato *anthocyanin reduced* (*are*) mutant has decreased levels of flavonol antioxidants in pollen and higher levels of reactive oxygen species (ROS). This mutant is hypersensitive to HT applied during pollen development and pollen tube growth, resulting in reduced pollen viability, pollen germination, and pollen tube elongation, suggesting that managing ROS levels is critical for pollen development and function under temperature stress ^61,62^.

In this study, we define the effects of HT on the pollen tube growth phase in a small set of tomato cultivars including the Heinz reference and others noted for fruit production at HT (Fig. 1A, Sup. Table S1). We show that application of acute HT exclusively during pollen tube growth (12 hours) is sufficient to reduce fruit biomass and seed number and that this detrimental effect is lessened in thermotolerant cultivars. We further show that pollen tubes from thermotolerant cultivars have enhanced pollen tube growth at HT *in vivo* and *in vitro*, suggesting that the capacity of thermotolerant cultivars to set fruit at HT is mediated by thermotolerant pollen tube growth. We used RNA-seq analysis of growing pollen tubes responding to HT to test hypotheses on the cellular basis of reproductive thermotolerance and define two response modes that distinguish thermotolerant cultivars: increased activation of stress response pathways and higher basal levels of growth promoting pathways the are repressed by heat stress.

Further, we identify increased expression of genes encoding key callose synthesis and flavonol synthesis enzymes as potential drivers of reproductive thermotolerance. Our work establishes the pollen tube growth phase as a tractable system for studying the impact of heat stress on cells and as a critical focal point for improving reproductive thermotolerance.

## Results

To determine whether cultivars that set fruit at HT show thermotolerant pollen tube growth, we searched the hundreds of available seed stocks at the UC Davis/C.M. Rick Tomato Genetics Resource Center and identified seven that were noted for fruit set at high temperature (Sup. Table S1). We used the Heinz reference and a set of three cultivars noted for thermotolerant fruit set so that we could perform multiple assays of pollen tube function and gene expression on each genotype: Tamaulipas, Malintka, and Nagcarlang are from the tropical coast of the Gulf of Mexico, Russia, and the Philippines, respectively. Each of these cultivars have been shown to produce fruits at high temperature in multiple field trials ^63,64,65–68^.

To determine the genetic relationships among the Heinz reference and the three cultivars noted for thermotolerant fruit set, we constructed a phylogeny that included our four test cultivars, a diverse set of 34 tomato cultivars, and one accession of *Solanum pimpinellifolium* to root the tree (Fig. 1A). Genome sequences were available for all ^42^ but Tamaulipas, so we performed short-read sequencing of the Tamaulipas genome (BioProject ID: PRJNA1128095). As previously reported ^42^, the Heinz reference was found in a clade that included other well known commercial cultivars (San Marzano, Brandywine, VF-36, Fig. 1A). Interestingly, Tamaulipas formed a clade with other cultivars identified on the Gulf Coast of Mexico/Central America, with Nagcarlang found in a related clade. Malintka was in a separate clade that included M-82 (widely used in tomato genetic analysis) ^69^. These results define the genetic relationships among our test cultivars, showing that Tamaulipas and Nagcarlang are relatively closely related within a group of semi-domesticated Central American varieties and equally unrelated to Heinz and Malintka, which are from a group of Euro-American agricultural cultivars.

### High temperature during the pollen tube growth phase reduces seed production; thermotolerant cultivars maintain relatively high absolute seed number

Seed production is directly dependent on pollen tubes delivering sperm cells to ovules for double fertilization. So, we determined whether HT applied during the pollen tube growth phase affected seed number (Fig. 1C). We grew Heinz, Tamaulipas, and Nagcarlang plants under optimal conditions, conducted hand self-pollinations, then incubated pollinated plants at either 25°C or 37°C for 12 hours before returning plants to optimal (25°C) growth conditions for 14 days (Fig. 1B, see methods). Applying HT only during the pollen tube growth phase significantly decreased seed production in Heinz and Nagcarlang (Fig. 1C, 1D). Consistent with Tamaulipas being the most thermotolerant line, HT did not reduce seed production in this cultivar. We identified a positive correlation between number of seeds and fruit mass (Sup. Fig. S1) in each of our test cultivars, further highlighting the importance of successful fertilization for tomato fruit production. These data suggest that HT applied only during the pollen tube growth phase reduces seed production, likely due to decreased fertilization, which could result in decreased fruit biomass production.

### High temperature during the pollen tube growth phase reduces fruit biomass

Applying HT for just 12 hours during the pollen tube growth phase significantly decreased fruit biomass for all cultivars analyzed (Fig. 1E). These results suggest that the pollen tube growth phase is highly susceptible to heat stress, and can significantly impact fruit yield. Heinz had the greatest temperature-dependent decrease in fruit biomass when heat stress was applied only during the pollen tube growth phase (less than half the mass of fruits produced by control pollinations, Fig. 1E, 1F). Tamaulipas and Nagcarlang maintained more fruit biomass when pollination occurred at HT compared with Heinz (>50% of control fruit weight, Fig. 1F).

### High temperature, applied only during the pollen tube growth phase, negatively affects pollen tube growth in the pistil

The reduction in seed number (Fig. 1C) suggests that applying HT to the pollen tube growth phase disrupts the ability of pollen tubes to reach ovules. There was no significant change in Tamaulipas seed set when the pollen tube growth phase was subjected to HT, suggesting that Tamaulipas pollen tube growth may be thermotolerant. We used two methods to test these hypotheses, and pollen development occurred under optimal conditions in both. First, Figure 2A outlines how we completely covered the stigma with an excess of pollen (profuse hand pollination), exposed growing pollen tubes to HT for 12 hours, and then returned them to optimal conditions for 12 hours before measuring how far the longest pollen tubes had extended (pollen tube front) in the style of the pistil (top of stigma to junction with the ovary). HT significantly reduced the ability of pollen tubes to grow the length of the style and reach the ovary in Heinz, Nagcarlang, and Malintka, but the effect of HT was not significant in Tamaulipas (Fig. 2B). Second, we conducted limited pollinations under control or HT so that we could measure the length of individual pollen tubes in pistils (Fig. 2C). The lengths of individual pollen tubes were also significantly reduced by HT in all cultivars (Fig. 2D). This effect was significantly greater for Heinz when compared to Malintka, Nagcarlang, or Tamaulipas when we analyzed the relative pollen tube growth in the pistil at 37°C relative to 25°C (Fig. 2E). Tamaulipas pollen tube growth was the most resistant to HT (Fig. 2D, 2E). Furthermore, while HT significantly decreased the fraction of pollen tubes from Heinz that reached the ovary, HT had no effect on the fraction of pollen tubes that reached the ovary for Malintka and Tamaulipas (Sup. Fig. S2). Our results suggest that HT reduces fruit biomass and seed set by inhibiting pollen tube growth in the pistil such that pollen tubes are unable to reach and fertilize the ovules and that Tamaulipas is largely resistant to these effects.

**Figure 2:**
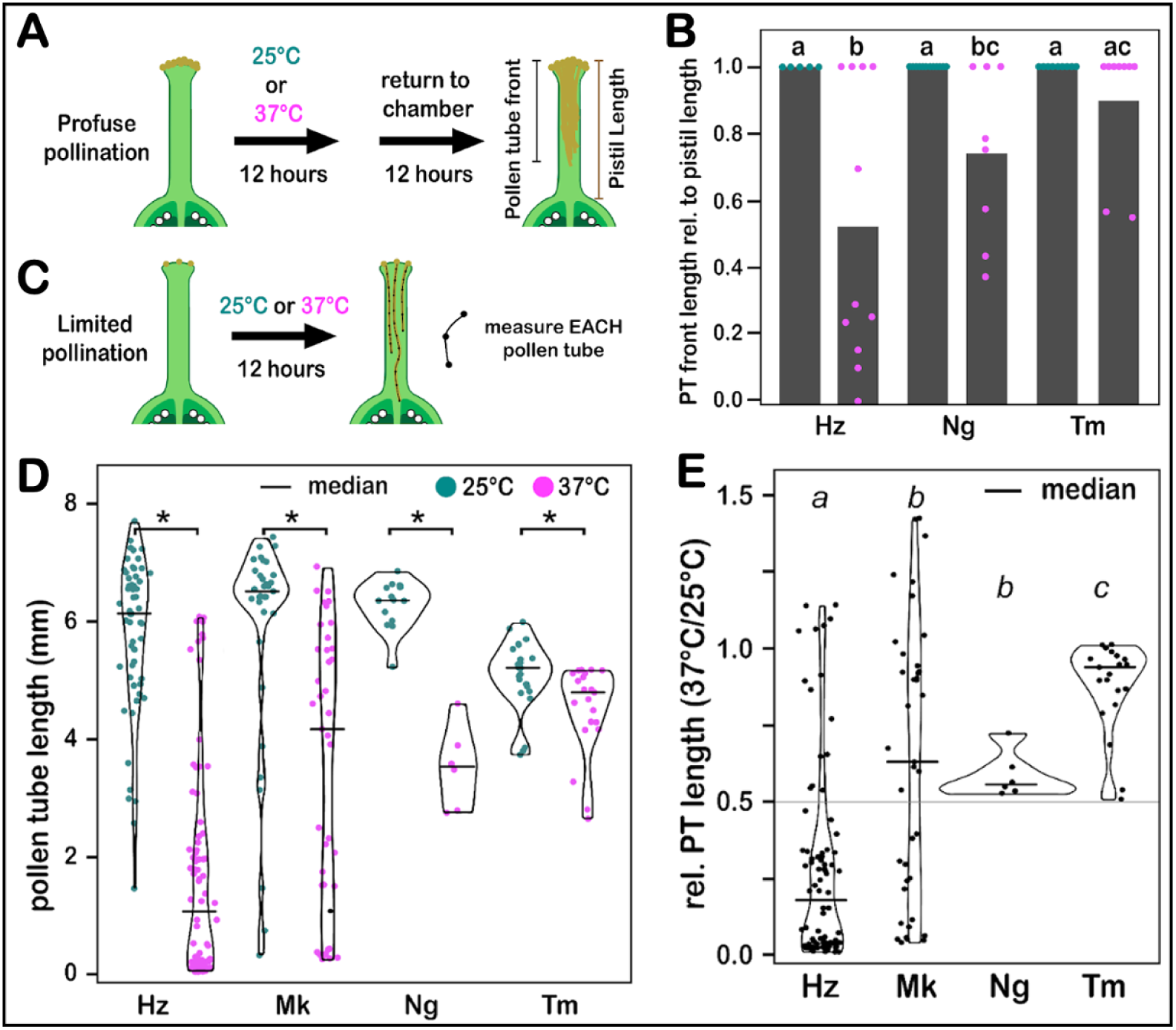
High temperature, applied only during the pollen tube growth phase, negatively affects the growth of pollen tubes in the pistil; TT cultivars show increased relative pollen tube growth *in vivo*. **(A)** Pollen grains that developed at 25°C were profusely hand pollinated onto emasculated pistils of the same cultivar and incubated for 12 hours at either 25°C (control) or 37°C (stress) during daytime, after which they were returned to their original cycle for 12 hours. The pollen tube front was measured relative to the pistil length. **(B)** Pollen tube group extension relative to the pistil length under control (dark green) and stress (magenta) conditions explained in A above. **(C)** Limited numbers of pollen grains that developed at 25°C were hand pollinated onto emasculated pistils of the same cultivar and incubated for 12 hours at either 25°C or 37°C, after which the length of each pollen tube in the pistil was measured. Pistils with less than 5 grains were used to measure the length of individual tubes. Pistils with more than 5 pollen grains were excluded from further analysis. **(D)** Pollen tube length inside the pistil after 12 hours at 25°C (dark green) or 37°C (magenta). **(E)** Relative pollen tube length in the pistil (37°C/daily, median value at 25°C). Statistical analysis was done using the Mann-Whitney *U* test for D (*ns, p >* 0.05; *\**, *p* ≤ 0.05), and using the Kruskal-Wallis test and Dunn’s test for B and E. Similar letters indicate no significant difference (*p* > 0.05) between groups from Dunn’s test. Hz, Heinz; Mk, Malintka; Ng, Nagcarlang; Tm, Tamaulipas.

### High temperature negatively affects the pollen tube growth *in vitro*

To begin to define how HT affects the growing pollen tube, we measured the lengths of pollen tubes generated from pollen grains that developed under optimal conditions and then were grown *in vitro* under control (28°C for 6 hours) or HT conditions (28°C for 3 hours, 37°C for 3 hours, Fig. 3A). Our goal was to ensure that pollen tubes had the opportunity to germinate and begin to extend pollen tubes under control conditions (28°C) so that we could evaluate the effect of HT on growing pollen tubes after 3 hours of stress treatment (Fig. 3A). We also measured pollen tube length at the three-hour time point to ensure that pollen tubes continued to grow during the stress treatment (Sup. Fig. S3A, S3B). Applying HT to pollen tubes extending *in vitro* resulted in a significant reduction in pollen tube length in all cultivars except for Tamaulipas, for which the decrease was not significant (6h Stress v. 6h Control, Fig. 3B, 3C, Sup. Fig. S3). Heinz experienced the largest absolute (Fig. 3C) and relative decrease in pollen tube length (Fig. 3D) after exposure to HT, while Tamaulipas had the highest relative pollen tube growth at HT compared to control (Fig. 3D). Our *in vitro* analysis suggests that pollen tube growth is significantly decreased by HT and that cultivars like Tamaulipas that set fruit at HT, may have genetic variation that allows them to maintain pollen tube extension under HT.

**Figure 3:**
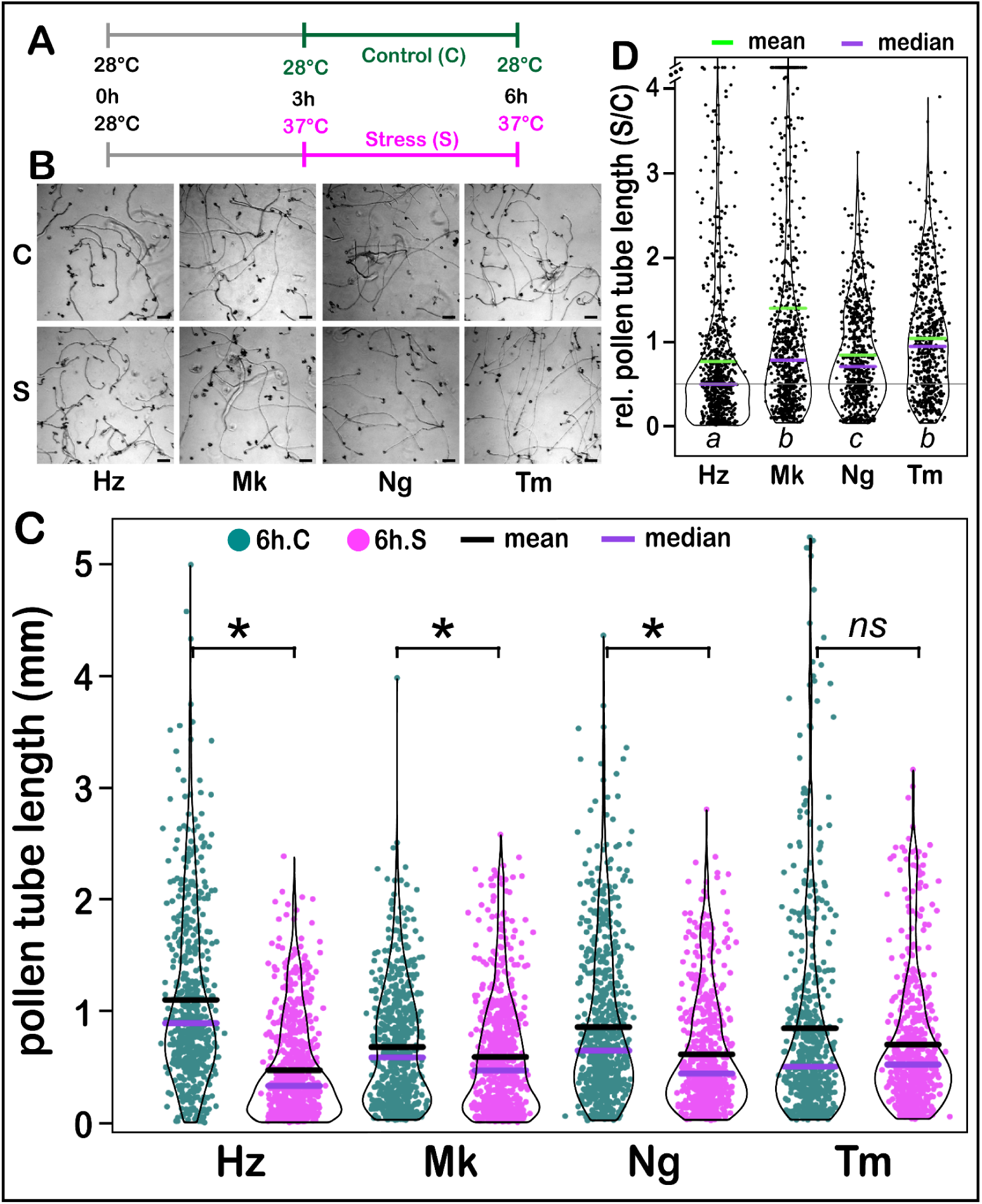
High temperature decreased pollen tube growth *in vitro*. **(A)** Pollen grains that developed at 25°C were collected from flowers 0 to 1 day after anthesis and incubated in liquid growth media for the indicated time and temperature. **(B)** Representative images of PT after growth, scale bar = 100 um. **(C)** *in vitro* PT length after 6 hours incubation at specified conditions. **(D)** Relative pollen tube length *in vitro* (Stress / daily median value of Control). Statistical analysis was done using the Mann-Whitney *U* test for C (*ns, p >* 0.05; *\**, *p* ≤ 0.05), and using the Kruskal-Wallis test and Dunn’s test for D. Similar letters indicate no significant difference (*p* > 0.05) between groups from Dunn’s test. Hz, Heinz; Mk, Malintka; Ng, Nagcarlang; Tm, Tamaulipas.

These experiments show that the Heinz reference and three cultivars noted for their ability to set fruit at high temperature display a range of sensitivity to HT applied during the pollen tube growth phase. Heinz was the most sensitive to temperature in all assays (Figs. 1-3), suggesting that pollen performance in the reference genotype is relatively thermosensitive. In contrast, Tamaulipas showed the highest degree of pollen thermotolerance in all assays (Figs. 1-3), with insignificant effects of HT for seed number (Fig. 1C) and pollen tube growth in vitro (Fig. 3C). Nagcarlang (closely related to Tamaulipas, Fig. 1A) and Malintka (equally unrelated to Heinz or Tamaulipas/Nagcarlang, Fig. 1A) generally outperformed the Heinz reference under HT (Figs. 1-3), but were not as thermotolerant as Tamaulipas for these parameters.

### Pollen tubes from thermotolerant cultivars mount a more robust heat stress response

Having established that the pollen tube growth phase is critically sensitive to HT (Fig. 1), and that cultivars of tomato that set fruit at high temperature display varying degrees of thermotolerant pollen tube growth in the pistil (Fig. 2) and *in vitro* (Fig. 3), we used RNA-seq to define the transcriptomes of pollen tubes growing under control or HT *in vitro* (using the same growth conditions as in Fig. 3A, control: 6 hours at 28°C, stress: 3 hours at 28°C and then 3 hours at 37°C). Our goal was to determine the molecular responses of pollen tubes in our test set of four cultivars to distinguish among three hypotheses for thermotolerance: 1) thermotolerant cultivars have a higher optimal temperature for pollen tube growth and thus do not respond to HT at the molecular level, 2) thermotolerant pollen tubes have enhanced induction of heat stress responses compared to that in thermosensitive cultivar pollen tubes, and 3) thermotolerant cultivar pollen tubes express stress responses under control conditions and are thus primed for a more effective response to heat stress.

Principal component analysis of our RNA-seq data set (two temperatures, four genotypes, three replicates) showed the majority of variation in the experiment was due to genotype rather than temperature, with Tamaulipas driving this component (PC1-axis, Fig. 4A). HT accounted for the second-highest component of the observed variation (PC2-axis, Fig. 4A). Analysis of the number of differentially expressed genes (DEGs, control v. stress for each cultivar, Fig. 4B, C, Sup. Tables, S5-8) indicated a modest pollen transcriptome response in all cultivars and that Heinz had the fewest responsive genes (103 DEGs, Fig. 4C), while Tamaulipas and Malintka had the most responsive genes (390 and 428 DEGs, respectively, Fig. 4C). Forty-two DEGs (the center of the Venn Diagram, Fig. 4C; core pollen tube HT-responsive DEGs) were common to all cultivars and showed similar directional expression changes across all (29 were upregulated in response to HT in all cultivars; 13 were downregulated in response to HT, Fig. 4D, Sup. Table S12). Thirteen of 29 core DEGs significantly increased by HT encode heat shock proteins (Fig. 4D; Sup. Table 12).

**Figure 4:**
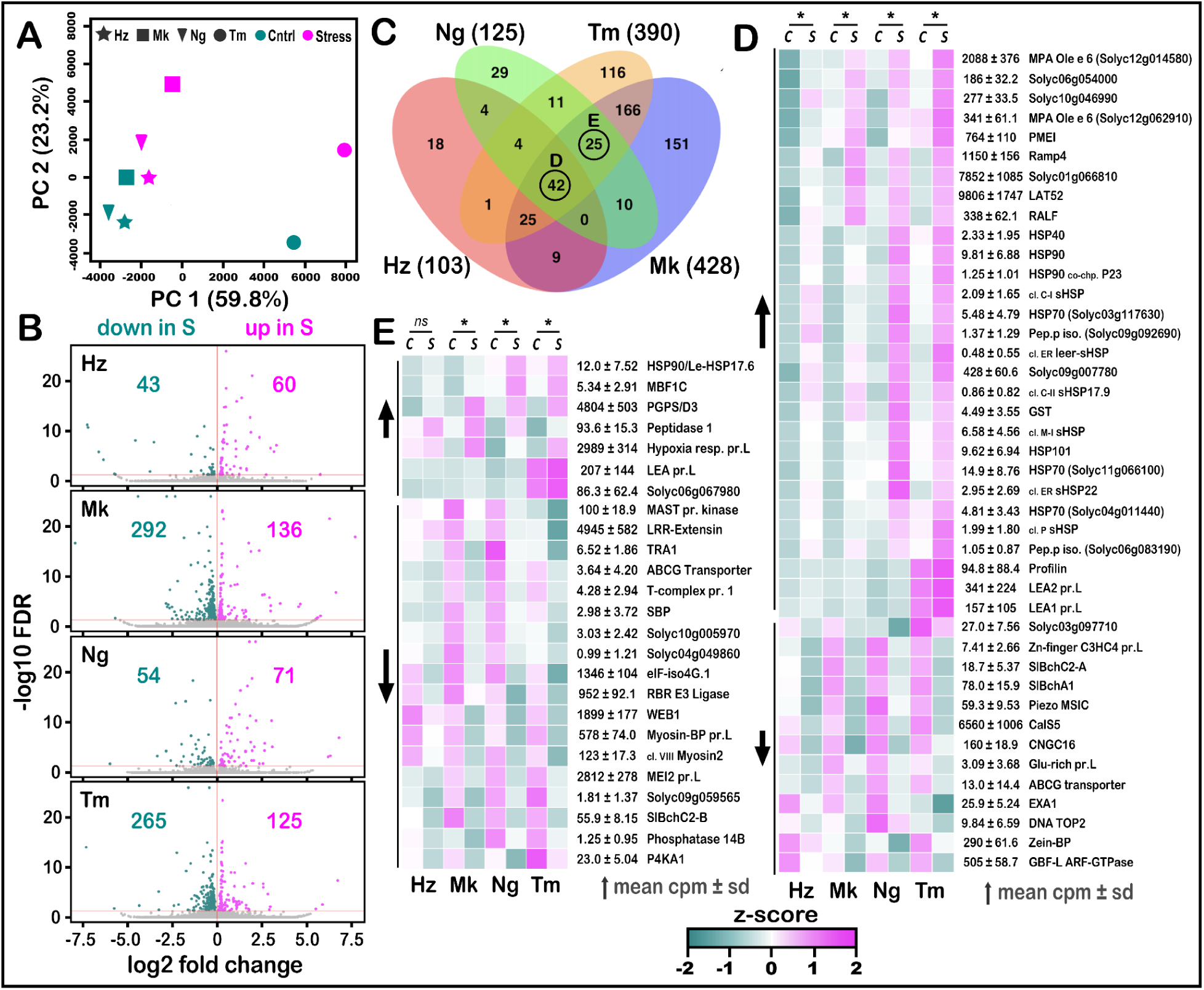
Growing pollen tubes exposed to high temperature mount a limited transcriptional response; TT cultivars show distinct and stronger responses. **(A)** The transcriptomes of pollen tubes grown *in vitro* for 6 hours, under the conditions highlighted in Figure 3A (28°C for 6 hours [control] and 28°C for 3 hours followed by 37°C for 3 hours [stress]), were resolved and analyzed using principal component analysis. **(B)** Scatter plots showing the number of genes whose transcript numbers were significantly downregulated or upregulated (DEGs, FDR < 0.05) in stress compared to control conditions for each tomato cultivar. **(C)** Venn Diagram of similarities among the DEGs shown in panel B; total number of genes with significantly altered expression in each cultivar are shown within brackets. Circles show core pollen tube HT-responsive DEGs (D) and core thermotolerant DEGs (E). **(D)** Heat map of 42 DEGs shared among all cultivars (core pollen tube HT-responsive DEGs). **(E)** Heat map of 25 DEGs shared by Mk, Ng, and Tm, but not Hz (core thermotolerant DEGs). Heat maps in D and E are shown as *Z*-scores, the number of standard deviations above (magenta), below (dark green), or at the mean (white) CPM across all treatments for each gene. *= significantly different when compared to control condition Hz, Heinz; Mk, Malintka; Ng, Nagcarlang; Tm, Tamaulipas; C, control (28°C, 6 hours); S, stress (28°C, 3 hours; 37°C, 3 hours).

Importantly, we also observed that induction was more pronounced in Nagcarlang and Tamaulipas for many of the core pollen tube HT-responsive DEGs (Fig. 4D, note higher levels on heat maps). We defined 25 DEGs that were unique to the three cultivars noted for thermotolerant fruit set (core thermotolerant DEGs, Fig. 4E). These DEGs, unlike the core pollen tube HT-responsive DEGs, were mainly decreased in response to HT. Among the small number of upregulated thermotolerant core DEGs was MBF1C, which has been identified as a HT response regulator that improves tolerance to stress and is independent of the heat shock factor (HSF) pathway ^74–80^. These data illustrate that both thermosensitive and thermotolerant cultivars have transcriptional responses to elevated temperatures, ruling out the hypothesis that thermotolerant cultivars have higher optimal temperatures. In contrast, we found support for the hypothesis that cultivars noted for thermotolerant fruit set have a more robust (more DEGs) heat stress response.

### High temperature leads to activation of a set of cellular and protein homeostasis responses that are shared across cultivars

To identify the molecular networks induced by HT in the growing pollen tube and to determine whether thermotolerant cultivars express different response networks, we performed gene-set enrichment analysis of functional categories on the set of 611 genes that were differentially expressed between control and HT in at least one cultivar (Fig. 4C, Sup. Tables S5-8). Despite the small number of DEGs that were observed, significant gene set associations were found in all eight annotations curated by STRING (Fig. 5, Sup. Figs. S5-10 ^81^). Response to temperature stimulus (heat stress) was among the biological processes activated in all cultivars (Fig. 5A). In addition, Nagcarlang significantly upregulated genes annotated as functioning in the response to oxidative stress, osmotic stress, and salt stress (Fig. 5A).

**Figure 5:**
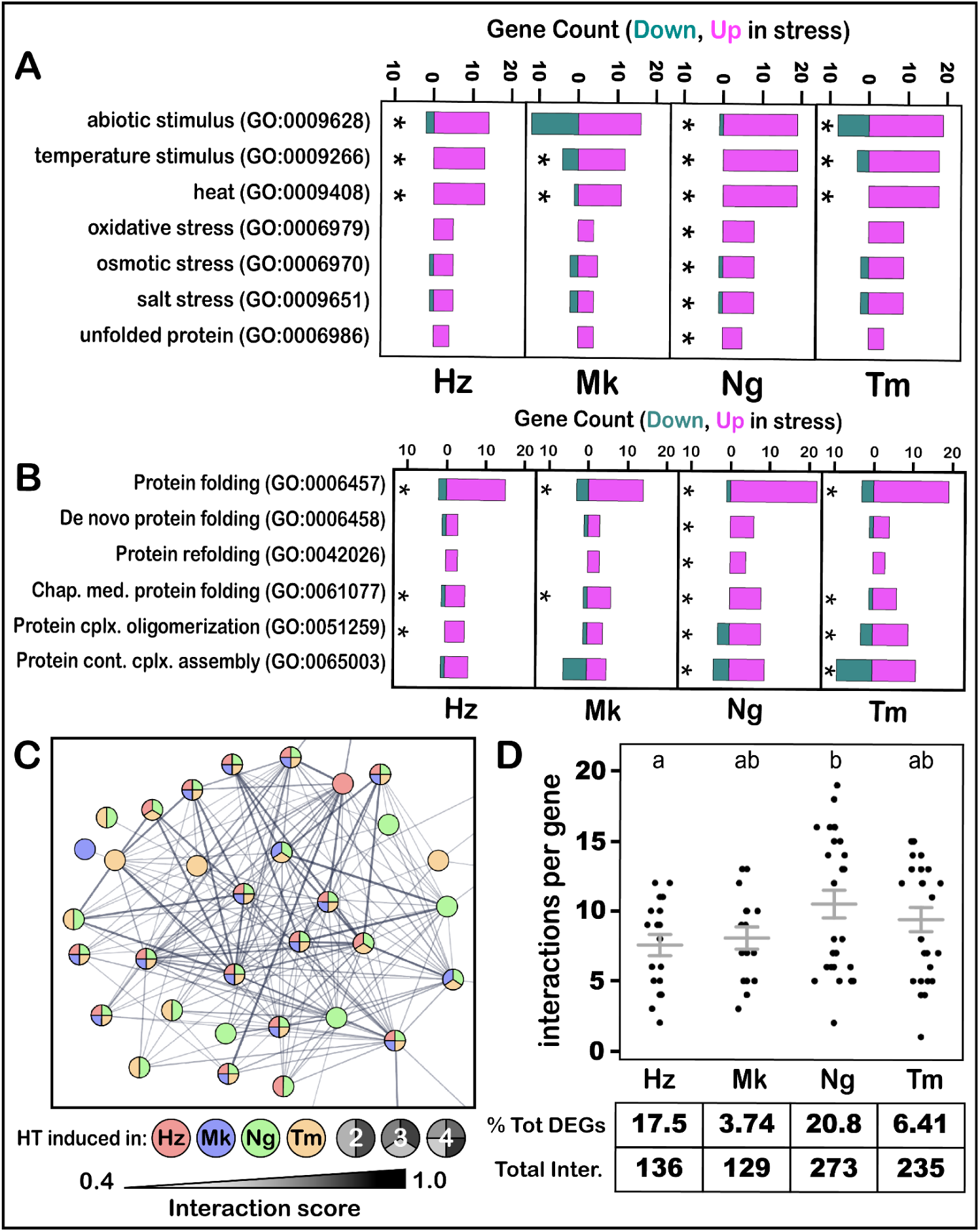
Pollen tubes of thermotolerant cultivars have a more robust response to high temperature. **(A)** Cell Responses, **(B)** Protein Homeostasis, **(C)** Visualization of the protein homeostasis network, each node (circle) is colored to show the cultivar(s) that use the node. **(D)** Quantification of interactions in C. Hz, Heinz; Mk, Malintka; Ng, Nagcarlang; Tm, Tamaulipas; stress (28°C, 3 hours; 37°C, 3 hours); control (28°C, 6 hours).

Other biological processes enriched in all cultivars were protein folding and chaperone-mediated protein folding (Fig. 5B) suggesting that HT leads to protein misfolding inside growing pollen tubes. We used STRING to define networks (based on published physical and functional interactions) among the DEGs ^81^. The most prominent network included genes involved in protein folding. We found that the size (Fig. 5C, number of nodes in each cultivar) and connectivity of this network (Fig. 5D, interactions per gene) varied across the four cultivars. Nagcarlang had more genes in the network (Fig. 5C, green and red) as well as significantly more interactions per gene compared to Heinz (Fig. 5D). These data indicate that the core responses to HT are the cellular response to heat and the unfolded protein response. The increased number of genes and network connections further support the hypothesis that the pollen tubes of thermotolerant cultivars (especially Nagcarlang and Tamaulipas) induce more robust response pathways compared to Heinz.

### Thermotolerant cultivars are transcriptionally primed to respond to high temperature

Finally, we asked whether our transcriptomic data provided evidence for priming of stress response pathways in pollen tubes of thermotolerant cultivars. We considered two forms of priming, 1) pre-induction: thermotolerant cultivars express higher basal levels of genes that are induced by HT in thermosensitive Heinz, 2) anti-repression: thermotolerant cultivars express higher basal levels of genes that are downregulated by HT and thus maintain higher levels under HT. We reasoned that either or both of these priming mechanisms would allow thermotolerant cultivars to maintain pollen tube growth under HT.

To determine whether cultivars noted for thermotolerant fruit set show pre-induction priming, we asked whether genes that significantly changed in thermosensitive Heinz exposed to HT (103 genes total, Fig. 4C) were also significantly different between thermotolerant cultivars and Heinz under control conditions. If pre-induction priming was occurring, we would expect to find that significant numbers of genes altered in Heinz under HT were already elevated under basal (control conditions) in the other cultivars, especially in Tamaulipas, which demonstrated thermotolerant pollen performance (Figs. 1-3). We identified 60 genes that were significantly upregulated and 43 genes that were significantly downregulated in Heinz exposed to HT (Fig. 4B, Fig. 6A top row). Many of the 60 genes upregulated in Heinz under exposure to HT, were found to be significantly higher in thermotolerant cultivars compared to Heinz under control conditions (Fig 6A, bottom three rows: Malintka, 21.7%, 13 genes; Nagcarlang, 26.7%, 16; Tamaulipas, 63.3%, 38 genes). The finding that nearly 2/3 genes upregulated in Heinz in response to HT were elevated in control conditions in Tamaulipas suggests that pre-induction priming of stress responses could be an important route to enhanced pollen performance under HT. In contrast, we identified very few genes that were upregulated in response to HT in Heinz that had significantly lower levels of expression in thermotolerant cultivars compared to Heinz under control conditions (0, 2, and 2 genes were ‘downregulated instead’). These data suggest significant pre-induction priming in thermotolerant cultivars for genes encoding chaperones and other protein folding factors, pollen growth factors, cell wall synthesis enzymes, cytoskeleton, and reactive oxygen species (ROS) scavengers (Fig 6B).

**Figure 6:**
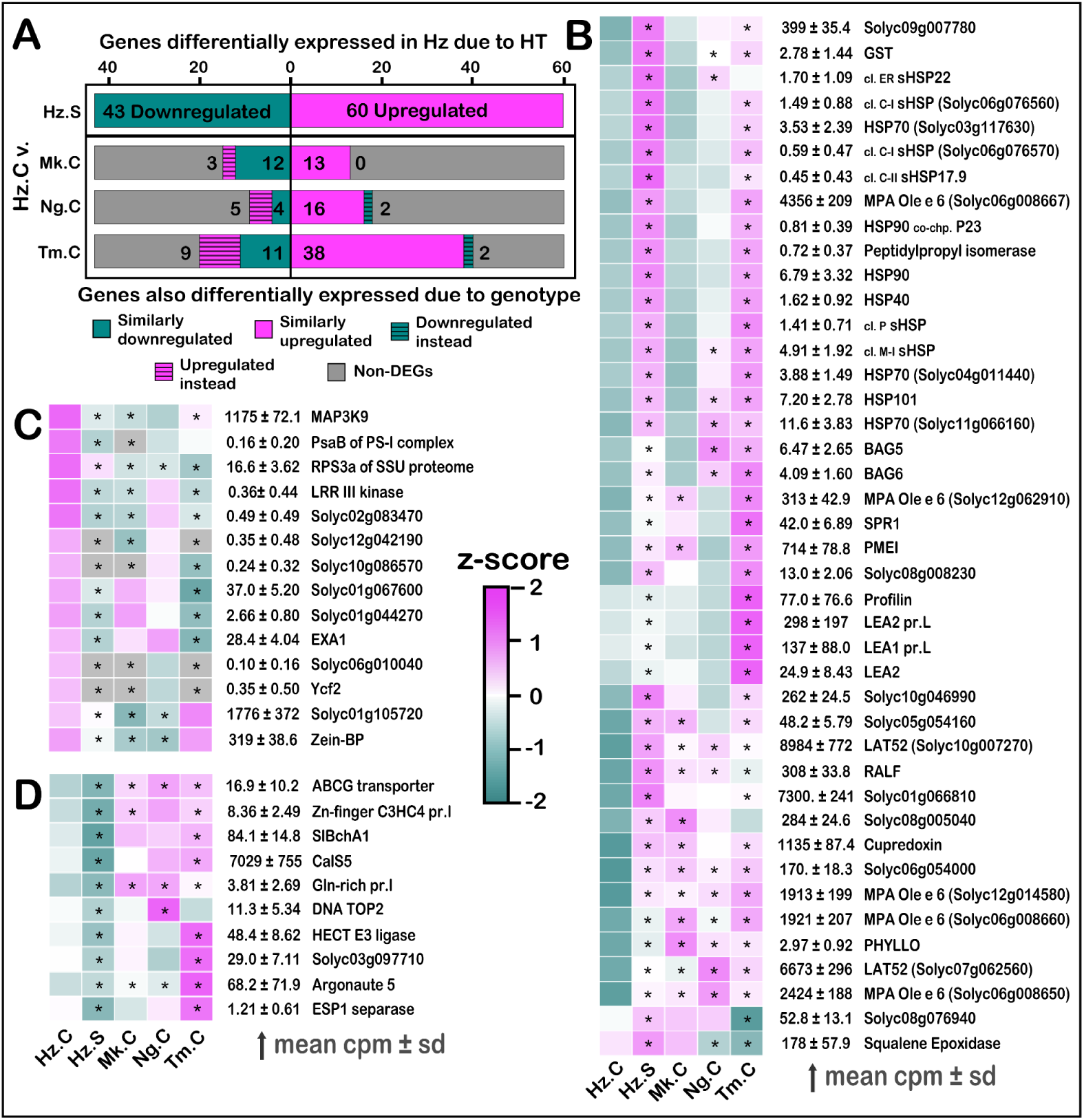
The thermotolerant pollen tube transcriptome is primed for the response to heat stress. **(A)** Genes that are upregulated or downregulated in Heinz in response to HT are represented in the top row; those same genes that are significantly upregulated/downregulated in another cultivar relative to Heinz are shown in the subsequent rows. **(B)** Genes that are induced by HT in Heinz and significantly higher in TT cultivars under control conditions. **(C)** Genes that are downregulated by HT in Heinz and significantly lower in TT cultivars under control conditions. **(D)** Genes that are downregulated by HT in Heinz and significantly higher in TT cultivars. **(B-D)** display *Z*-scores, the number of standard deviations above (magenta), below (dark green), or at the mean (white) CPM across indicated treatments for each gene. Hz, Heinz; Mk, Malintka; Ng, Nagcarlang; Tm, Tamaulipas; S, Stress (28°C, 3 hours; 37°C, 3 hours); C, Control (28°C, 6 hours).

On the other hand, of the 43 genes with transcripts significantly decreased in response to HT in Heinz, we found fewer genes that were significantly lower in thermotolerant cultivars compared to Heinz under control conditions (12, 4, 11 genes similarly downregulated, Fig. 6A, 6C) and we found similar numbers of genes that were significantly higher in thermotolerant cultivars compared to Heinz under control conditions (upregulated instead, n = 3, 5, 9, Fig. 6A, 6D). These genes show anti-repression priming—they are downregulated by HT in Heinz but have higher basal levels in thermotolerant cultivars—and include CALLOSE SYNTHASE 5 (CalS5), a critical enzyme for pollen tube cell wall formation ^82^, and the ortholog of Arabidopsis SPIRRIG (SlBchA1, an endosomal trafficking factor that has also been shown to be essential for tip growth in root hairs ^83,84^, Fig. 6D). These data provided support for our hypothesis that thermotolerant pollen tubes are primed for a more effective response to heat stress, particularly in Tamaulipas. These observations suggest that pre-induction priming and anti-repression priming are important modes of thermotolerance in growing pollen tubes.

### Tamaulipas pollen tubes maintain lower levels of ROS under control and heat stress conditions

Response to oxidative stress (Fig. 5A) and ROS (H_2_O_2_, Sup. Fig. S10) were among the biological processes found to be significantly induced by HT in our RNA-seq analysis and ROS levels have been shown to be critical for pollen tube performance. ROS levels at the lower extreme correlate with lack of pollen germination ^85^ and ROS levels increase with elevated temperature and at the higher extreme are associated with pollen tube growth defects ^61,62^. We tested whether thermotolerance can be explained by altered ROS levels in thermotolerant cultivars following HT. The ROS response to HT is known to be rapid ^86,87^, so we allowed pollen tubes to grow for one hour under optimal conditions (28°C) before applying HT (37°C) for one hour and measuring ROS levels in the first 30 µm of the actively extending pollen tube tip. Control pollen were grown at 28°C for two hours (Fig. 7A).

**Figure 7:**
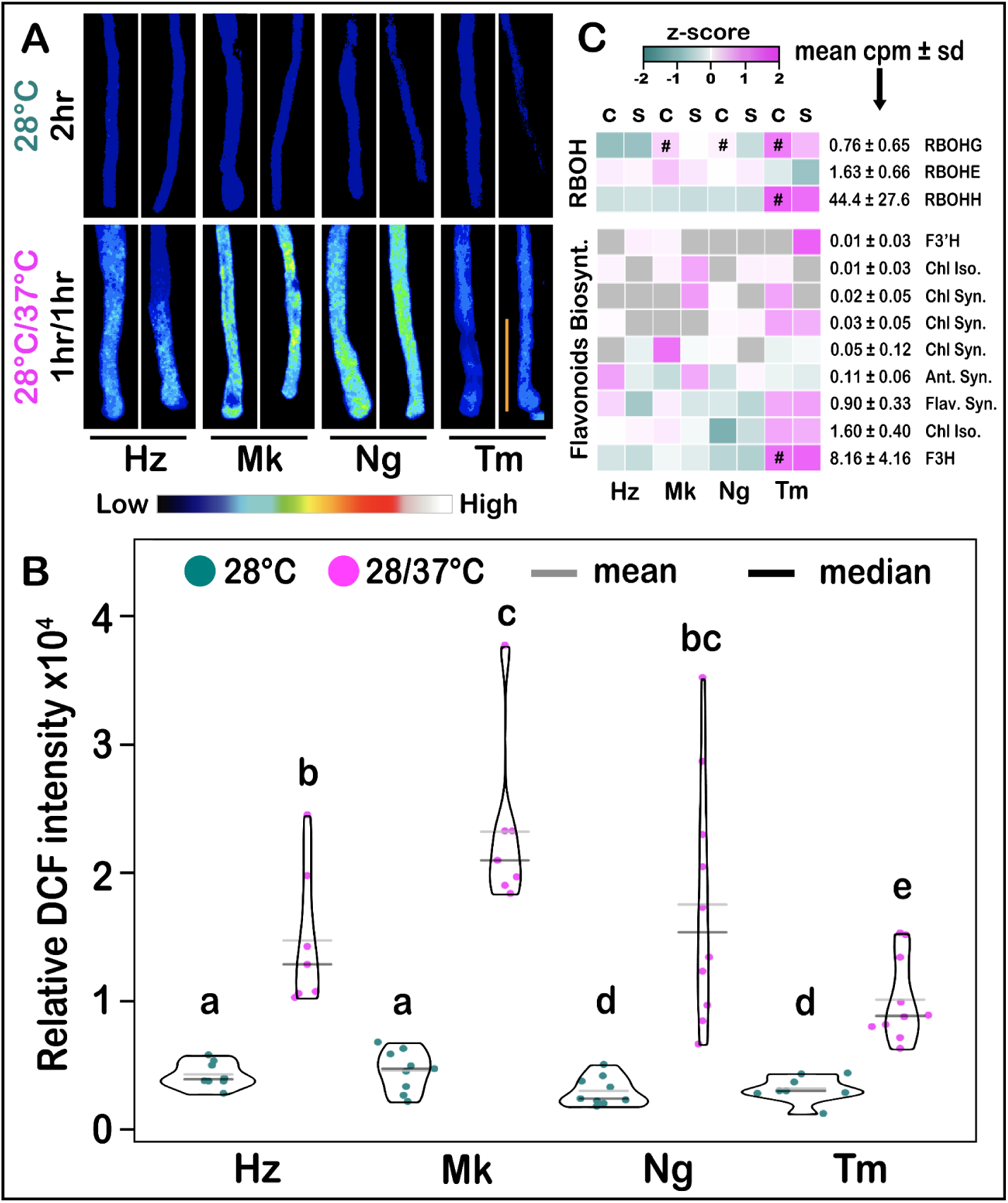
High Temperature increases ROS levels of growing pollen tubes. **(A)** Representative image of DCF fluorescence in pollen tubes germinated *in vitro* for 1 hour at 28°C then exposed for 1 more hour to 28°C or 37°C. The orange bar represents 30 µm (distance from the tip used to quantify relative DCF intensities). **(B)** Relative ROS levels measured as relative DCF fluorescence. Statistical analysis was done using the Kruskal-Wallis test and Dunn’s test. Similar letters indicate no significant difference (*p* > 0.05) between groups from Dunn’s test. **(C)** Heatmaps of gene families known to be involved in ROS synthesis (top) and scavenging (bottom). C, Control (28°C, 6 hours); S, Stress (28°C, 3 hours; 37°C, 3 hours). * = Stress significantly different from Control within cultivar. # = significant compared to Heinz at 28°C. Heatmaps display *Z*-scores, the number of standard deviations above (magenta), below (dark green), or at the mean (white) counts per million (CPM) across all treatments for each gene. Hz, Heinz; Mk, Malintka; Ng, Nagcarlang; Tm, Tamaulipas.

All cultivars had comparable ROS levels at 28°C with similarly low variation (Fig. 7A, 7B). However, HT significantly increased ROS levels in growing pollen tube tips of all cultivars (Fig. 7A, 7B), with the variation in pollen tube ROS also increasing, especially in Nagcarlang. This suggests that subpopulations of pollen tubes respond to HT by producing different levels of ROS (Fig. 7B). Nagcarlang and Malintka showed the largest increases in ROS in response to HT (Fig. 7C). Tamaulipas had significantly lower levels of ROS at HT than all other cultivars. It was notable that Tamaulipas, the most thermotolerant cultivar we analyzed, had lower ROS levels under control and HT and less variation under HT compared with other cultivars (Fig. 7B).

Analysis of the transcripts of genes required for ROS synthesis (respiratory burst oxidase homolog H, RBOH gene family) and scavenging (antioxidant flavonols) indicated that none are induced by HT in growing pollen tubes (Fig. 7C). This is consistent with the proposal that ROS levels increase rapidly under HT by post-transcriptional processes in growing pollen tubes. On the other hand, when we compared the levels of expression in control samples, we found that Tamaulipas pollen tubes have significantly higher transcript levels of the synthesis gene RBOHH and the gene encoding the flavonol biosynthetic enzyme F3H (the enzyme that is nonfunctional in the *are* mutant ^61,62^) relative to Heinz, providing examples of priming. These data, in combination with the observation that Tamaulipas is better able to maintain ROS levels under control and HT (Fig. 7A, B), suggest that Tamaulipas transcriptome is tuned to manage elevations of ROS levels that accompany pollen tube growth under HT.

### Tamaulipas maintains higher levels of callose synthesis under HT

HT results in repression of genes associated with growth in animal cells ^88–90^. The hypothesis is that the heat stress response includes a growth pause to maintain viability during stress. So, we sought to identify pollen tube growth pathways directly affected by HT that could mediate thermotolerance. The highly abundant Cals5 transcript decreases significantly in response to HT in all cultivars tested, a striking observation as pollen tube growth is inextricably linked to cell wall synthesis (CalS5, mean CPM 6560, Fig. 4D). This finding led to the hypothesis that repression of CalS5 is part of the pollen tube heat stress response and that thermotolerant cultivars, especially Tamaulipas, may have altered expression of callose synthases that enhances pollen tube growth under HT (Fig. 2, Fig. 3). To begin to test these hypotheses, we defined how pollen tube-expressed callose synthases respond to HT (Fig. 8A). Four additional callose synthase genes were also significantly decreased in response to HT (indicated by an asterisk, Fig. 8A), suggesting coordinated downregulation of members of this gene family. Importantly, we also found that Tamaulipas had significantly higher transcript levels under control conditions for four callose synthases (including the most abundant member of the family, CalS5) compared to Heinz after HT (#, Fig. 8A). This latter finding is consistent with anti-repression priming of callose synthesis in Tamaulipas, which may contribute to its ability to maintain pollen tube growth under HT (Fig. 2, Fig. 3).

**Figure 8:**
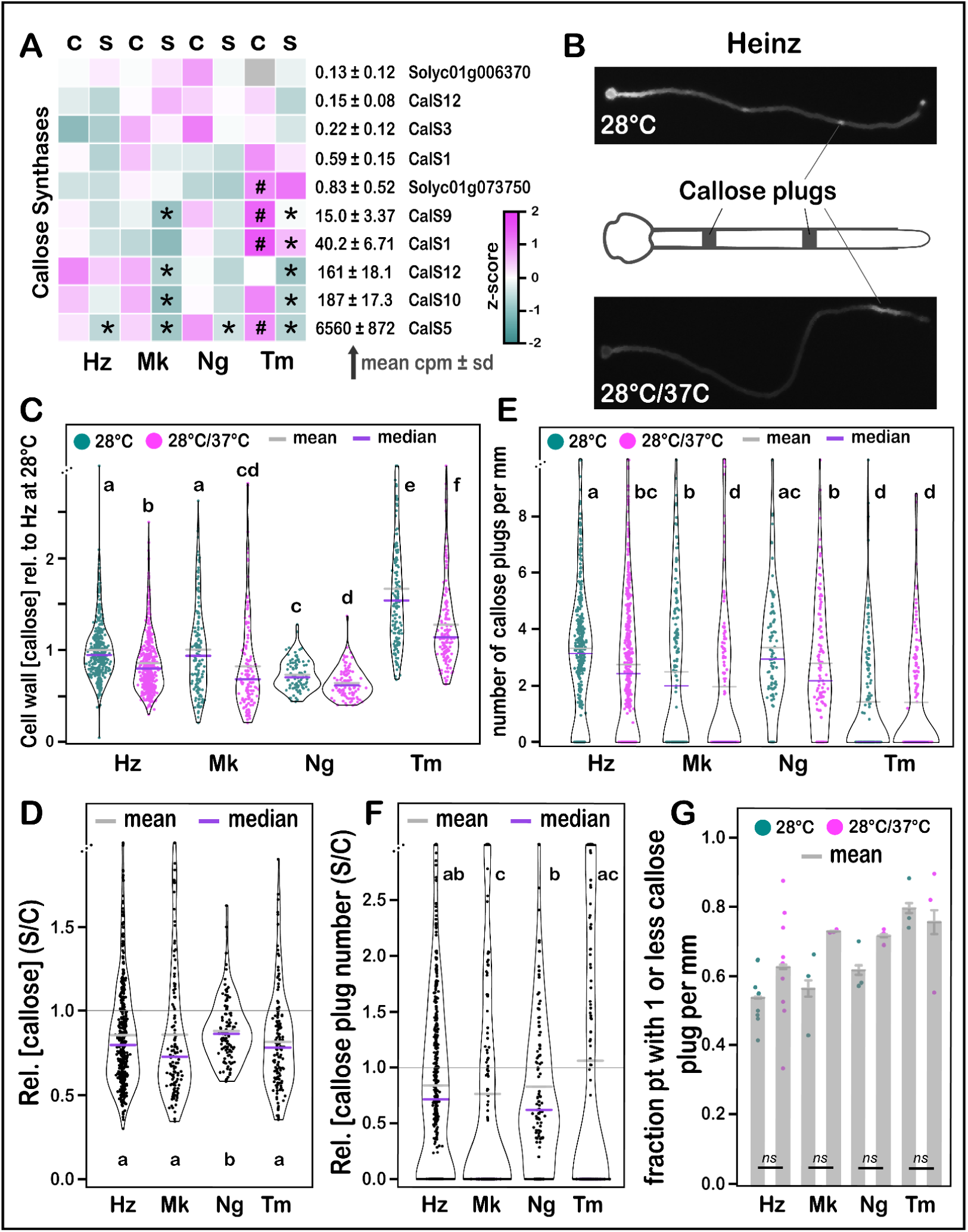
High temperature reduces callose deposition in the cell wall and the density of callose plugs. **(A)** Heatmap of annotated and pollen tube expressed (TPM > 0.1) callose synthase genes. * = significant compared to the same cultivar’s 28°C treatment; # = significant compared to Heinz at 28°C; *Z*-scores, the number of standard deviations above (magenta), below (dark green), or at the mean (white) CPM across all treatments for each gene. **(B)** Representative image of aniline blue stained Heinz pollen, grown under stress (S, 28°C, 3 hours; 37°C, 3 hours) or control (C, 28°C, 6 hours) conditions (as in 3A), showing callose plugs and callose levels along the pollen tube length. **(C)** The concentration of callose in the pollen tube cell wall normalized to the daily median value of Heinz at 28°C. **(D)** Relative cell wall callose concentration within each cultivar (stress/control). **(E)** The density of callose plugs relative to that of Heinz at 28°C. **(F)** Relative density of callose plugs within each cultivar (stress/control). **(G)** Fraction of pollen tubes for which we recorded one or less callose plug per millimeter, bars represent standard error. Statistical analysis was done using the Kruskal-Wallis test and Dunn’s test for C-G. Normalization to Heinz at 28°C in E and F were done using the daily mean instead of median to avoid dividing by 0; normalization for C and D was done using the daily median for Heinz at 28°C. Similar letters indicate no significant difference (*p* > 0.05) between groups from Dunn’s test. Hz, Heinz; Mk, sMalintka; Ng, Nagcarlang; Tm, Tamaulipas; S, Stress (28°C, 3 hours; 37°C, 3 hours); C, Control (28°C, 6 hours).

To determine if HT-mediated repression of callose synthase transcript abundance resulted in decreases in pollen tube callose concentration and to test for an altered response in thermotolerant cultivars, we analyzed aniline blue-stained pollen tubes subjected to the same control and HT conditions used in our RNA-seq analysis (Fig. 3A, Fig. 8B). HT led to a significant reduction in callose content in pollen tubes of all cultivars (Fig. 8C, D). Interestingly, and consistent with callose synthase gene expression changes in RNA-seq analysis, Tamaulipas pollen tubes had the highest baseline callose content at 28°C and maintained a significantly higher level of callose concentration than all genotypes under HT (Fig. 8C). These data indicate that HT leads to a decrease in callose in the pollen tube cell wall, which could directly result in reduced growth/extension (Fig. 2, Fig. 3), and that Tamaulipas is able to maintain growth under HT by expressing higher levels of pollen tube callose synthases.

In addition to synthesizing a major pollen tube cell wall component, callose synthase is also essential for production of callose plugs ^82^; thin (∼2 µm) depositions of callose that span the diameter of the pollen tube and make a physical barrier between the actively extending tip and the inactive distal region of the pollen tube (Fig. 8B, ^91^). Callose plug deposition is proposed to modulate pollen tube growth rate by regulating the turgor pressure (the driving force for extension) of the actively extending tip ^92,93^. We analyzed patterns of callose plug deposition to determine if HT alters this important component of pollen tube growth. HT led to a significant reduction in callose plug density (callose plugs per millimeter) in Heinz and Nagcarlang (Fig. 8E), but not in Tamaulipas (Fig. 8E-F). Tamaulipas had the lowest callose plug density under control conditions but maintained a similar density under HT (Fig. 8E-F). Furthermore, all cultivars, except Tamaulipas, experienced a slight increase in the number of pollen tubes that had one or less callose plug per millimeter (Fig. 8G). These data suggest that higher levels of callose and the ability to maintain patterns of callose plug deposition have contributed to pollen tube thermotolerance in Tamaulipas.

## Discussion

### The pollen tube growth phase is critical for seed and fruit production under HT; cultivars that maintain fruit production in hot growing seasons have thermotolerant pollen tubes

We found that HT applied for only 12 hours during the pollen tube growth phase was sufficient to significantly reduce fruit biomass in all cultivars tested (Fig. 1B, C, D), but that Tamaulipas was resistant to the HT-induced reduction in seed number observed in other cultivars (Fig. 1E, F). These findings suggest that Tamaulipas produces fruit under HT because its pollen tubes are able to grow under HT and fertilize ovules to produce seeds, initiating successful fruit development. The relationship between seed number and fruit weight is complex and parthenocarpic tomato cultivars produce fruits with few or no seeds ^54,94^. However, in the cultivars we’ve studied, fruit biomass is correlated with seed number and seedless fruits have low fruit weight (Sup. Fig. S1). These findings suggest two paths to thermotolerant fruit production: 1) a high basal seed number that decreases under HT but stays above a threshold (Nagcarlang, Fig. 1E), or 2) the ability to maintain near optimal levels of seed set under HT (Tamaulipas, Fig. 1E, F).

Our hypothesis was that Tamaulipas (and other thermotolerant cultivars) are able to produce seeds under HT due to enhanced pollen tube growth. In experiments conducted in the pistil (Fig. 2), we found that HT decreases pollen tube growth in all cultivars, but the effect was less pronounced in Tamaulipas and Nagcarlang (Fig. 2B, D, E). *In vitro* pollen tube growth experiments yielded similar results. Pollen tube growth is repressed by HT in all cultivars, but the impact of heat stress was more significant in Heinz (Fig. 3C). Our phenotypic analysis shows that enhanced pollen tube growth under HT is an important contributor to thermotolerant fruit production.

### Thermotolerant pollen tube growth is mediated by a combination of enhanced induction and priming of stress response pathways

Our analysis of how the pollen tube transcriptome responded to HT showed that temperature was a key driver of the variation observed in our RNA-seq data set (Fig. 4A) and that all cultivars induced a heat stress response (Fig. 4B, 4D, 5A). These observations lead us to reject the hypothesis that thermotolerant cultivars have higher optimal temperatures. In contrast, the data supported the hypothesis that thermotolerant cultivars have an enhanced heat stress response. We observed greater numbers of DEGs (Fig. 4B) and higher levels of induction (Fig. 4D) in thermotolerant cultivars, and significant induction/repression of genes that don’t change in thermosensitive Heinz (Fig. 4E). In particular, we found more nodes and more connections in the protein homeostasis networks induced by HT in thermotolerant cultivars (Fig. 5B, C, D). A more robust heat stress response was particularly evident in Tamaulipas (hyper induction of HSPs, Fig. 4D) and Nagcarlang (more robust response network, Fig. 5D).

Tamaulipas, the cultivar that best maintains fruit set, seed set, and pollen tube growth under heat stress, suggests that priming is an important mode of thermotolerance. Almost two thirds of the genes induced by heat stress in Heinz had significantly higher levels under control conditions in Tamaulipas relative to Heinz (Fig. 6A, 6B). This enrichment shows a powerful example of pre-induction priming. Tamaulipas has high basal levels of genes that thermosensitive Heinz induces following exposure to HT. These genes include many encoding proteins that maintain proper folding of other proteins at HT (Fig. 5B) such as heat shock protein chaperones, BAG family chaperones, and protein folding catalysts Peptidylpropyl Isomerases (HSP, BAG, Peptidyl isomerase, Fig 6B). In addition, we saw evidence for anti-repression priming (expression of genes at higher basal levels that are repressed by HT), thus maintaining expression above a threshold under HT (Fig. 6A, 6D).

### Tamaulipas is primed for expression of genes involved in ROS synthesis and scavenging

ROS are known to play key roles in pollen performance and it has come to be appreciated that maintaining optimal ROS levels is critical - pollen failure is associated with low or high levels of ROS ^95,96^. Specific ROS may play direct roles in spatio-temporal modulation of pollen tube cell wall physical properties or activity of enzymes that synthesize the cell wall or maintain tip growth and pollen tube extension. Hydroxyl radicals (·OH) have been proposed to sever pollen cell wall polymers to promote pollen tube germination and flexibility of the extending tip, while hydrogen peroxide (H_2_O_2_) limits tube extension by increasing cross-linking of polymers and increasing rigidity ^97^ and can increase the oxidation of cysteines in proteins to affect their function, including enzymes and transcription factors ^98^. Loss of function of pollen tube tip-localized RBOH (*rbohh*, *rbohj* double mutants) results in pollen tubes that burst prematurely in Arabidopsis, and apoplastic ROS produced by tip-localized RBOH has been hypothesized to be required for pollen tube integrity ^99,100^. In tobacco, decreases in pollen tube extension due to reduction of expression of RBOH could be restored by addition of H_2_O_2_, suggesting that H_2_O_2_ (a product of the combined activity of RBOH and SOD) is the ROS critical for pollen tube integrity ^97,101^.

We found that HT caused significant increases in ROS levels in growing pollen tubes in all of the cultivars we analyzed (Fig. 7A, 7B), which is in keeping with the well established relationship between heat and oxidative stress ^87^. However, Tamaulipas had the lowest levels of pollen tube ROS under both control (except for Nagcarlang) and HT conditions, which suggests that ROS management could contribute to pollen tube thermotolerance.

Cellular ROS levels are set by the interplay of respiration, enzymatic ROS production from RBOHs, and scavenging by anit-oxidant metabolites and enzymatic antioxidants ^86,102^. To gain insight into how Tamaulipas maintains ROS homeostasis, we analyzed expression of the RBOH gene family and genes required for flavonol biosynthesis (Fig. 7C). Flavonols are powerful antioxidants and it has been shown that pollen of tomato *are* mutants, which lack an important flavonol synthesis enzyme (F3H), have lower levels of pollen flavonols, higher levels of pollen tube ROS, and decreased pollen viability, germination, and tube growth under HT ^61,62^. Moreover, pollen performance under HT can be improved by adding flavonols to growth media, by inhibiting ROS synthesis, or by overexpression of the F3H enzyme ^62^.

Tamaulipas shows significantly higher levels of the *F3H* transcript, which encodes a key enzyme in flavonol biosynthesis, under control and stress conditions than Heinz (Fig. 7C), with trends toward higher levels of transcripts of other genes encoding enzymes in this pathway. Thus, Tamaulipas may be better able to modulate ROS levels than the other cultivars we’ve analyzed. However, Tamaulipas also showed higher basal levels of the major pollen tube RBOH (RBOHH, Fig. 7C) and of SOD (Supplementary Fig. S11). Future work will be required to determine how these (and other) critical determinants of cellular ROS levels are regulated to achieve enhanced ROS homeostasis in thermotolerant Tamaulipas.

### Pathways controlling pollen tube growth and cell wall synthesis are repressed by HT; Tamaulipas is primed to maintain growth

Repression of growth is part of the heat stress response in many systems ^88–90^ and our data suggests pollen tube extension is also repressed by heat stress (Fig. 2, Fig. 3). Analysis of the pollen tube transcriptome highlighted multiple genes associated with pollen tube growth that were repressed in response to HT (Fig. 4D, 4E). The most abundantly expressed callose synthase (CalS5) was significantly decreased in response to HT in all cultivars (Fig. 4E, Fig. 8A). An interesting next step would be to test whether CalS5 is a target for transcriptional repression by heat shock factors and other transcription factors that respond to heat stress in the growing pollen tube (Fig. 4E). Repression of CalS5 is expected to decrease growth because callose is both a major component of the pollen tube cell wall and critical for formation of callose plugs ^82,92^. Our data suggest that repression of CalS5 following heat stress is a critical driver of the decrease in pollen tube growth we observe in the pistil (Fig. 2) and *in vitro* (Fig. 3).

We also identified two homologs of Arabidopsis SPIRRIG with transcripts that significantly decrease in response to heat stress in all cultivars (SlBchA1 and SlBchC2-A, Fig. 4D, Sup. Fig. S4) and a third member of this family (SlBchC2-B, Sup. Fig. S4) that also decreases in response to HT in all cultivars, with significant decreases in thermotolerant cultivars (Fig. 4E). SPIRRIG is a member of the deeply conserved Beige and Chediak Higashi (BEACH) domain-containing protein family ^103^ and was shown to be essential for extension of tip-growing root hair cells. Localization of SPIRRIG to the root hair tip has been correlated with root hair extension rate ^83^. BEACH domain-containing proteins are large and have multiple functional domains ^104^. In plant cells, these proteins are critical regulators of actin dynamics required for endosomal trafficking in tip-growing root hairs ^83^ and have been implicated in the response to salt stress via formation of processing bodies (mRNP granules that facilitated mRNA turnover during periods of abiotic stress) ^105^. Thus, these proteins can integrate two important heat stress responses in pollen tubes: repression of tip growth and degradation of mRNA to bolster protein homeostasis ^106,107^. We propose that the tomato homologs of SPIRRIG expressed in pollen tubes (SlBchA1, SlBchC2-A, SlBchC2-B, Fig. 4D, E, Sup. Fig. S4) are targets for repression of pollen tube extension during heat stress.

Thermotolerant cultivars, especially Tamaulipas, accumulate higher transcript levels of CalS5 and the tomato ortholog of SPIRRIG (SlBchA1) under control conditions (Fig. 6D) and maintain higher transcript levels under HT (Fig. 4E). In the case of CalS5, we have shown that these changes in gene expression are associated with increased pollen tube callose content under HT in pollen tubes of Tamaulipas (Fig. 8C) and propose that anti-repression priming of CalS5 contributes to Tamaulipas pollen tube thermotolerance (Fig. 2, Fig. 3). Thermotolerant Tamaulipas shows significant anti-repression priming of SlBchA1 (Fig 6D). These data suggest that increased basal expression levels of these factors that promote tip growth and modulate stress responses can help achieve thermotolerance in pollen tubes. These proposals will be the focus of future experiments aimed at determining whether increased CalS5 or SlBchaA1 expression is required for Tamaulipas thermotolerance.

### Callose deposition is critical for thermotolerant pollen tube growth

Turgor pressure is thought to be the driving force for pollen tube extension ^92,93^ and the pollen tube cell wall is constructed to maintain structural integrity via a rigid shank (distal part of the pollen tube) as the endomembrane system delivers material (membrane, cell wall polymers, proteins) to the more fluid tip ^108^. The force of turgor pressure must be spatio-temporally balanced by the pollen tube cell wall to maintain rapid extension without structural failure. We propose that this balance is disrupted by HT and that thermotolerant cultivars express mechanisms that enhance this balance under stress. We found that HT results in lower callose levels in pollen tubes (Fig. 8C). Treatment of pollen tubes with callose degrading enzymes showed that callose is required to maintain cell wall integrity ^109^, perhaps because pollen tubes with reduced callose are unable to resist turgor pressure. Thus, Tamaulipas pollen tubes may be thermotolerant because they maintain higher levels of callose under HT (Fig. 8C).

Callose plugs partition the extending pollen tube into smaller pressure units; a decrease in the number of callose plugs is expected to lead to an increase in volume and a decrease in turgor pressure in each unit ^92^. We found that HT reduces the number of callose plugs in pollen tubes of Heinz, Malintka, and Nagcarlang, but not of Tamaulipas which was already primed for higher pressure unit volumes (lower callose plug density under control, Fig. 8E, F). The greatest decrease was in Nagcarlang (Fig. 8F), suggesting that Nagcarlang may achieve pollen tube thermotolerance via modulation of callose plug deposition at different temperatures. Tamaulipas, however, may achieve pollen tube thermotolerance via pre-induction priming to absorb any additional turgor pressure due to HT. Our RNA-seq data suggest that callose synthase transcript abundance, as well as cell wall callose content, are heat stress-responsive (Fig. 8A). Future investigations should focus on whether the activity of this plasma membrane-localized enzyme is altered in response to changes in temperature and pressure to optimize callose plug deposition for optimal cellular extension.

### Prospects for future development of this experimental system

By conducting experiments that limit application of heat stress to the pollen tube growth phase, we have shown that this brief window of the life cycle is critically important for fruit and seed production at HT (Fig. 1). We have also demonstrated that Tamaulipas, which like Nagcarlang and Malintka is a cultivar noted for its ability to maintain higher fruit production during exceptionally hot growing seasons ^63,64,68,70,72,73^, is able to fertilize seeds and set fruit when stress is only applied during the pollen tube growth phase (Fig. 1) and has enhanced pollen tube performance under HT in all of the assays we performed (Fig. 2, Fig. 3). Moreover, we tested multiple hypotheses for the molecular basis of pollen tube cellular thermotolerance and found that a combination of enhanced induction and priming of stress responses (Fig. 4-8) are associated with thermotolerant pollen tube growth under HT (Fig. 2, Fig. 3). Importantly, we have identified components of the pollen tube stress response (Fig. 4D, Fig. 5), ROS homeostasis (Fig. 7), and callose synthesis and deposition (Fig. 8) as major focal points for reproductive thermotolerance. Our work provides resources that will accelerate efforts to breed and engineer reproductive resilience in crops in a changing climate.

Pollen tubes offer an attractive model system for understanding how cells respond to heat stress and to define mechanisms of cellular thermotolerance that can be translated into improved crop resilience. Pollen is amenable to integration of single cell data that address cellular physiology/performance (Fig. 2, Fig. 3), genome-wide patterns of transcription (Fig. 4), and biochemical responses (Fig. 7A, Fig. 8B). Future work will be aimed at enhancing this experimental system by incorporating live-cell imaging and genomic analysis of earlier time points in the pollen tube growth process, which will allow us to determine how heat stress affects pollen germination, extension, and maintenance of cell wall integrity. Exploring the role that the pistil plays in supporting pollen tube growth in thermotolerant cultivars is critical also, as we work to identify gene variants that drive thermotolerance.

## STAR Methods

### Tomato Cultivars

Seeds of Heinz 1706 - BG (LA4345), Tamaulipas (LA1994), Malintka 101 (LA3120), and Nagcarlang (LA2661) were obtained from the Tomato Genetics Resource Center, UC Davis (Sup. Table S1). Seeds and cuttings were used to raise plants in the greenhouse and controlled growth chambers.

### Construction of a phylogeny of tomato Cultivars

Sequence FASTQ files were obtained from the NCBI Sequence Read Archive (Sup. Table S18) from 37 accessions originally sequenced previously ^42,45^, and a whole-genome sequence set from Tamaulipas generated as part of this study. Reads were mapped using BWA (v0.7.17-r1188 ^110^) to the SL4.0 reference genome ^111^ (www.solgenomics.net) using default settings. VCF files were imputed using bcftools (v1.12, Li 2011) with depth ≥ 2 and including both invariant and variant sites:

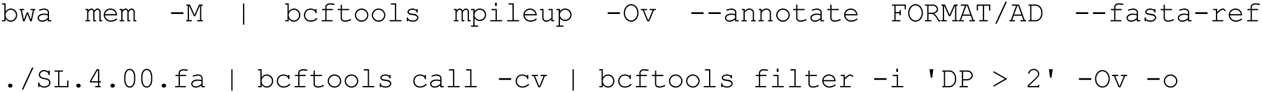

VCFs were converted to MVF files and merged into a single reference-anchored alignment using MVFtools ^112^. A FASTA concatenated alignment of 14,546,831 bp was generated from all sites with at least 34 out of 38 accessions represented. The alignment included 12.1% gaps and 80.4% of sites were invariant. A phylogeny of the 39 genomes (38 samples and the SL4.0 Heinz reference) was inferred using raxml-ng ^113^ using the GTR+F0+G4m model and all other settings were default values.

### Whole Genome Sequencing of Tamaulipas

Tamaulipas seedlings were germinated on soil for one week and then subjected to a 48 hour dark treatment before DNA was extracted from above ground portions of seedlings following a previously published method ^45^. Genomic DNA was sequenced using Illumina High-Seq; data are available at the short-read archive (https://www.ncbi.nlm.nih.gov/sra) using accession number: PRJNA1128095 or SRX25048037.

### Tomato plant growth

The University of Arizona (analysis of seed/fruit production, Fig. 1; analysis of pollen tube growth in the pistil, Fig. 2; analysis of callose content in pollen tubes, Fig. 8) plants were grown in Conviron growth chambers maintained at either 16-h-day/8-h-night, maintained at 25°C and 15°C, respectively or13-h-day/11-h-night, maintained at 25°C and 17°C, respectively. The light intensity was set to a maximum of 400 µmoles per sq. cm. Plants were watered regularly with a 0.5x nutrient solution of either ICL Peters Professional 5-11-26 Hydroponic Special or as reported previously ^114^.

Brown University (*in vitro* pollen tube growth, Fig. 3; RNA-seq analysis, Fig. 4) plants were grown under a 13-h-day/11-h-night greenhouse cycle in two-gallon pots and fertilized with Israel Chemicals Limited Peters Professional 5-11-26 Hydroponic Special. The temperatures were weather-dependent and fluctuated between day temperatures of 21°C to 28°C, and night temperatures between 16.5°C and 18°C. Fruits and stems were trimmed for maximum flower production. The plants were grown either from rooted cuttings of self-fertilized tomato lines or by seed germination. Tomato seedlings were germinated from potting soil planted seeds under standard greenhouse conditions (16-h-day/8-h-night cycle, 25 °C/18 °C day/night temperature) in a controlled growth chamber environment for two weeks before being transferred to a two-gallon pot.

Wake Forest University (analysis of ROS levels, Figure 7A) plants were grown in a greenhouse with day temperatures set to 28°C and night temperatures set to 21°C.

### Flower staging, pollen collection, and flower pollination

Flowers were staged based on the opening of buds (Sup. Fig. S12A) and the yellowness of the anther cone (Sup. Fig. S12B). Pollen samples were collected by vibration of the anther cone of Stage +1 flowers and grown in liquid media or hand-pollinated onto Stage 0 pistils that were emasculated approximately 24 hours prior (at Stage -1).

### Analysis of fruit and seed production (Fig. 1)

The morning before pollination and temperature stress, the oldest buds in an inflorescence with yellowing corollas were tagged and emasculated by removing the anther cone and leaving behind the exposed pistil (https://tgrc.ucdavis.edu/pollinating). This ensured that the pistils were not inadvertently self-pollinated. Approximately 24 hours after emasculation, the pistils reached maturity and were receptive to pollen and supported pollen tube growth, which was confirmed with previous aniline blue staining experiments. For each cultivar, pollen from at least two opened flowers (Stage +1 or +2 obtained from at least one plant in each condition) was collected by touching excised anther cones with a hand-held electric toothbrush modified to hold a collection tube, pooled onto a sterilized surface, and used to self-pollinate emasculated flowers. The pistils were profusely pollinated by completely coating the stigma with pollen. Following the pollination of all emasculated flowers, the plants were incubated in temperature chambers that were set to either 25°C (control) or 37°C (heat stress) for 12 hours. After 12 hours of exposure to either condition, all plants were returned to the control growth conditions (25°C/light for 13 hours and 17°C/dark for 11 hours). After 14 days, the tagged emasculated flowers were evaluated for the fruit set, which was determined by the presence or absence of a fruit. If a fruit formed, it was weighed and the mass was recorded as the 14-day fruit mass. Seeds were extracted with tweezers from the fruits and the number was recorded as seed set.

### Aniline blue staining assay of *in vivo* pollen tube growth (Method 1, Fig. 2A)

As described for fruit mass/seed yield experiments, all pistils were emasculated, profusely pollinated by hand, and exposed to either control conditions (25℃) or heat stress conditions (37℃) for 12 hours. After this exposure, the plants were returned to control conditions (25℃/light for 13 hours and 17℃/dark for 11 hours). The aniline blue staining procedure was inspired by protocols from the Dr. Patricia Bedinger’s laboratory and used previously ^115^. Approximately 24 hours after pollination, the hand-pollinated pistils were excised and placed stigma-side down into a labeled 0.8 mL tube containing 300 μL of fixative solution (3:1 95% EtOH: glacial acetic acid) and incubated for a minimum of 16 hours. After at least 12 hours, the fixative solution was removed, and 300 μL of 5 M NaOH was added to soften the tissue. The pistils remained in the softening solution overnight, but not more than 24 hours. The 5 M NaOH solution was removed after 12-24 hours, and the pistils were gently washed three times with 300 μL of distilled water. After the third wash was removed, 300 μL of prepared aniline blue solution (1:20 stock 0.1 mg/mL aniline blue dissolved in distilled water:0.1 M K_2_HPO_4_, pH 10 buffer) was added to each tube. The pistils with the aniline blue solution were placed in the dark at room temperature for at least 24 hours. The stained pistils were placed on a microscope slide, approximately 10 μL of mounting solution (1:3 50% glycerol:stock 0.1 mg/mL aniline blue dissolved in distilled water) was added, and a coverslip was gently placed overtop the pistil and pressed to flatten the ovary. The slides were observed with an Axiovert microscope under UV irradiation conditions.

### Aniline blue staining assay of *in vivo* pollen tube growth (Method 2, Fig. 2C)

Flower staging, emasculation, and pollen collection were performed as previously described for the fruit mass/seed yield experiments. 24 hours after emasculation, stigmas of emasculated pistils were gently dabbed in isolated pollen on a sterile surface in such a way that only a few pollen grains were deposited on the stigma. Pollinated pistils were incubated in the control or heat stress conditions for 12 hours. Pistils were excised and an aniline blue assay of pollinated pistils was performed as described ^116^. Stained pistils were mounted and observed using epifluorescence microscopy on a Zeiss Axiovert 100 fluorescent microscope.

### *In vivo* pollen tube length measurements (Fig. 2)

To image aniline blue stained pistils, the UV light was filtered through recommended filter sets (UV-1A, UV-2A, and UV-2B), which have an excitation and emission spectrum, of 370 and 509 nm, respectively. 5X magnified Images were captured using a Retiga Exi camera and the Metamorph software. Pollen tube length measurements were made using the publicly available NIH Image J software. Pixel to length conversion was calculated using a reference micrometer image captured under the same settings. For profusely pollinated assay (Fig. 2A, B), the front of the pollen tubes (area where the majority of the pollen tubes stop) and the pistil (area from the top of the stigma to the junction of the style and ovary) lengths were measured using Image J software. The ratio of the pollen tube front length to the pistil length was calculated and recorded. For the limited pollination assay (Fig. 2C-E), pollen tube length measurements were performed only in those pollinated pistils that contained ≤ 5 pollen grains on the stigma (Sup. Fig. S13) by tracing the pollen tube using the freehand tool in NIH Image J and the length of each trace were measured as described for profuse pollination experiments.

### Pollen growth medium

Pollen growth medium (PGM) was made as described ^115^: 24% (w/v) polyethylene glycol (PEG) 4000, 0.01% (w/v) boric acid, 2% (w/v) Suc, 20 mM MES buffer, pH 6.0, 3 mM Ca(NO3)2·4H2O, 0.02% (w/v) MgSO4·7H2O, and 1 mM KNO3.

### Analysis of pollen tube growth *in vitro*

Brown University (Fig. 3, Fig. 4): On the morning of the experiment, pollen was collected into 0.5 mL conical tubes. Dry pollen samples were homogenized in 250 µl PGM, diluted 1:6 into three new Eppendorf tubes (50 μL pollen solution in 250 μL fresh pollen growth medium), and grown for 3 hours at 28°C. After 3 hours of growth, 1 tube was collected for brightfield imaging on poly-D-lysine coated slides. The second tube was left at 28°C and the last tube moved to 37°C for 3 hours. After the last 3 hours, the two remaining tubes were collected for imaging as explained above. Pollen tube length was measured using the segmented perimeter tools on ImageJ. A pollen grain was considered germinated if the length of the pollen tube was greater than the diameter of the pollen grain. The limits of the pollen tube were defined as the length between the edge of the germinating pollen grain and the tip of the pollen tube. Pollen tubes were only measured if their limits were clearly defined in a single image or a combination of multiple images. In each measured image, all defined germinated pollen grains were analyzed. Pollen tubes were imaged at the highest concentration of pollen in the dish in which pollen tube limits were clearly defined. At least 100 pollen tubes were measured in each sample.

### Analysis of pollen tube growth *in vitro* for aniline blue staining

The University of Arizona (Fig. 8): On the day of the experiment, open flowers (stage +1) were excised from Heinz and either Malintka, Tamaulipas or Nagcarlang plants, and pollen from one flower of each cultivar was used in each replicate. 3 replicates were performed in total for Malintka, Tamaulipas and Nagcarlang with Heinz used as a control cultivar in each replicate. Pollen was collected by cutting off the tip of the anther cone and a modified electric toothbrush was used to buzz the pollen from the anther cone into microcentrifuge tubes containing the PGM and mixed well. PGM-containing pollen was then used to prepare a serial dilution and placed in two 24-well glass-bottomed plates previously coated overnight with 100 μg/mL poly-D-lysine dissolved in water and then thoroughly rinsed with excess water. One plate was incubated at 28°C for 6 hours while the other was incubated at 28°C for 3 hours followed by 37°C for another 3 hours.

### Pollen tube RNA extraction

Pollen tubes were grown *in vitro* as described above and collected in 0.75 mL Eppendorf tubes by centrifugation and flash frozen in liquid nitrogen. Pollen tube samples were placed in frozen metal blocks (liquid nitrogen in a styrofoam bath) and were ground using a plastic pestle for 1-2 minutes. The pollen was kept frozen during the grinding process. RNA was isolated following the RNeasy Plant Mini Kit protocol. After grinding the pollen, Qiagen RLT buffer was added to the Eppendorf tube without removing the pestle. The pestle was only removed after all material was washed from the tip. After isolation, all RNA samples were treated with DNase. To each 30 μL of RNA solution, 3 μL of 10x Turbo DNase buffer was added and mixed followed by 1 μL of DNase. After incubation at 37°C for 20 minutes, 6 μL of DNase inactivation agent was added to each sample. Samples were centrifuged at 10,000g and the supernatant was moved to a separate Eppendorf tube for storage at -80°C.

### RNA Sequencing Analysis

RNA-seq data are provided on the short-read archive (https://www.ncbi.nlm.nih.gov/sra, SRP252265). RNA samples were sequenced by Illumina High-Seq and the resulting reads were mapped to the tomato reference genome (SL4.0, ITAG4.1) following NextFlow’s RNA-seq pipeline (https://nf-co.re/rnaseq). The resulting salmon.merged.gene_counts were analyzed for differentially expressed genes with EdgeR ^117^. The principal component analysis (Fig. 4A) was calculated using the average transcripts per million value for each gene across three replicates for each cultivar at each temperature. Gene set enrichment analysis was performed using STRING version 11.5 ^81^. Protein sequences from ITAG 4.1 were first custom annotated against the more than 14000 organisms in the string database (using the annotation feature of STRING) to create a custom protein network. The annotated genome is accessible and usable in the string database under <u>STRG0093MZF</u>. The list of differentially expressed genes, obtained within each cultivar due to HT, were then compared against this custom genome to generate lists of overrepresented pathways. For all analyses, a false discovery rate (FDR) of 0.05 was used as a cutoff and no fold change cutoff was applied due to the limited number of differentially expressed genes obtained.

### *In vitro* callose measurements using an aniline blue analysis

At the end of incubation, plates containing *in-vitro* grown pollen tubes were stained with aniline blue by pipetting out the PGM from each well and first replacing it with fixative solution (see above). After a 30-minute incubation at room temperature, the fixative solution was removed and replaced with 50% ethanol solution and incubated for another 10 minutes at room temperature. The 50% ethanol was then removed and replaced with decolorized aniline blue solution (0.1% (w/v) aniline blue in 108 mM K_3_PO_4_. pH ∼11) as prepared according to ^118^. The plates were then incubated for 30 minutes in the dark before imaging. Pollen tubes were visualized on a Zeiss Axiovert 100 microscope and 5 images covering different areas of each well were obtained. Exposure time (750 ms) remained consistent across all trials. Heinz was always used a reference to normalize day-to-day variation in detected fluorescence. Images were processed in Python using <u>FluoroQuant</u>, an application we developed to quantify pixel intensities–above a set background pixel intensity–from boxes added along the length of particles to quantify. The script is available at https://github.com/souonkap/FluoroQuant. For each measurement, a background pixel intensity of 10 was used. The pollen grain was excluded in each trace, as the fluorescence intensity in the grain would reflect callose deposition in the flower prior to heat stress experiment. For each pollen tube measured, we recorded the pollen tube length (in mm), the number of callose plugs, and the mean gray value (total gray value of all selected pixels divided by the number of pixels selected) normalized to that of Heinz at 28°C to compare average callose deposition.

### *In vitro* reactive oxygen species measurements

Pollen grains were collected from flowers at anthesis and were placed in 300 µL PGM in 24 well plates with glass bottoms for 1h at 28°C to allow germination and initiation of pollen tube growth. All cultivars were analyzed in parallel during each experiment. One set of samples was transferred to 37°C for 1h for temperature stress, while control samples remained at 28°C for the additional 1h. A 500 µM stock of 2’-7’-dichlorodihydrofluorescein diacetate (CM-H2DCFDA) (Thermo-Fisher) was prepared in DMSO and added to pollen to achieve a 5 μM working concentration. CM-H2DCFDA was added to wells 20 min before the end of incubation. All samples were imaged immediately after treatment. DCF fluorescence was visualized using a Zeiss 880 laser scanning confocal microscope (LSCM) using the 488 nm laser and an emission spectrum of 490-606 nm. Laser settings were uniform for all samples and all experiments. Laser power of 1% and gain of 980 were used. The DCF fluorescence intensity was quantified in FIJI.

### Statistical analysis

Unless otherwise stated, all statistical analyses reported were done using non-parametric statistical approaches (Fig. 1, Fig. 2, Fig. 3, Fig. 7B, C, Fig. 8C-G). To improve our ability to account for daily variation in experimental conditions, control and stress samples within each cultivar were always run in parallel and obtained from a single batch of pollen grains collected from flowers right before the start of each experimental trial. Furthermore, before statistical analysis, each raw measurement (V_r_) made at either control or stress temperatures was normalized (V_n_) to the same day/trial median (or daily mean when stated) measurement (V_dm_) observed for that cultivar at control temperature (V_n_ = V_r_/V_dm_). We then used a Kruskal-Wallis test to determine if significant variation existed in the entire experiment (all cultivars, temperatures, trials, and time points included). To define if HT had a statistical effect on the performance of a particular cultivar, we performed an analysis of medians using the Mann-Whitney *U* test on normalized temperature-specific distributions (Stress vs. Control). To determine if the cultivars performed differently from one another, we performed a Kruskal-Wallis test followed by Dunn’s test using only the normalized distributions at HT when the cultivar trials were obtained independently of one another. When the trials of all genotypes were done in parallel or when all trials included a genotype control (e.g., Heinz used in all independent runs), normalized values were obtained by dividing each data point by the daily median (or mean when stated) of the control genotype at the control temperature (Heinz at the lower temperature). Statistical significance was then obtained using a combination of Kruskal-Wallis and Dunn’s, or Mann-Whitney *U* tests. The statistical analysis of our RNA-seq data was determined using the EdgeR package of bioconductor^117^ and returned raw *p*-values were adjusted using the false discovery rate method of the internal p.adjust() function of R. All downstream analyses were done using the false discovery rate of 0.05 as a significance cutoff. Gene set enrichment analysis statistics were performed using the STRING database v11.5^81^.

## Acknowledgements

This project was funded by a grant from the US NSF (IOS-1939255, Co-PIs MAJ, RP, JBP, GKM, AEL) with additional support from USDA/NIFA grants (2020-67013-30907, Co-PIs GKM, MAJ) and (2024-67012-41882 to MFA), and NIH (5R35GM139609, PI AEL). We thank Sherry Warner (Brown University), Nick Vasquez (Brown University), and Chieri Kubota (University of Arizona) for expertise in growing and maintaining tomato plants. We thank our colleague, Alison DeLong and the Brown University undergraduates (Danielle Alvarez, David Barrera, Grace Cinderella, Emma Corcoran, Austin Draycott, Emma Herold, Arielle Johnson, Alex Markes, Tu Pham, and Maria Rodriguez) who contributed to the early development of many of the protocols and concepts used in this work, especially Ryan Chaffee who participated in early analysis of the RNA-seq data. We also acknowledge generous support for undergraduate research from the Brown University UTRA/SPRINT program and the ASPB SURF Fellowship (BS); from a Brown University Presidential graduate Fellowship (SOY); and from a Raman Fellowship for Post-Doctoral Research by the University Grants Commission of India (MP). Seeds were obtained from the UC Davis/C.M. Rick Tomato Genetics Resource Center, Department of Plant Sciences, University of California, Davis, CA 95616.

## Author contributions

S.V.Y.O, M.P., K.P., E.J. M.F.A. B.S. R.A.A. conducted the experiments and developed methods; S.V.Y.O, S.E.S, R.R, A.F.H, J.B.P, A.E.L analyzed data; S.V.Y.O, S.E.S, G.K.M., R.P., M.A.J. designed the experiments; S.V.Y.O, R.P., M.A.J. wrote the manuscript with input from all authors.

## Abbreviation

HT: high temperature

