## Supplementary material for "Enhanced pollen tube performance at high temperature contributes to thermotolerant fruit production in tomato": supplemetal materials

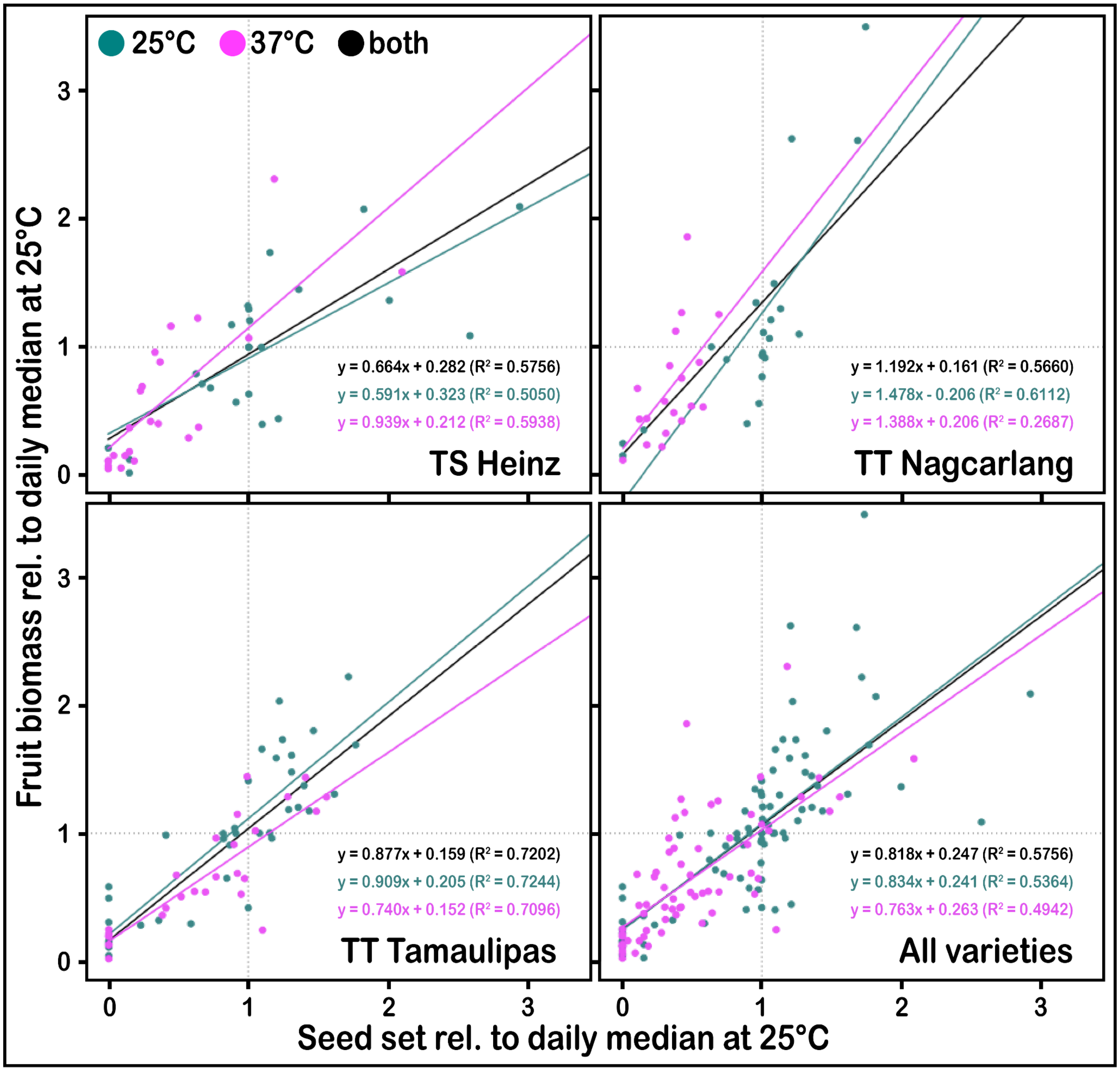


**Supplemental Figure S1. Correlation between fruit biomass and seed number in tomato at both optimal and high temperatures.** The relative biomass of each fruit (obtained under conditions highlighted in Figure 1A) was plotted against the relative seed set of the same fruit. All correlations were obtained using linear regression and returned positive correlations that were all significant ($p<0.05$). The $R^2$ values as well as the regression coefficients of each best fit line are reported.

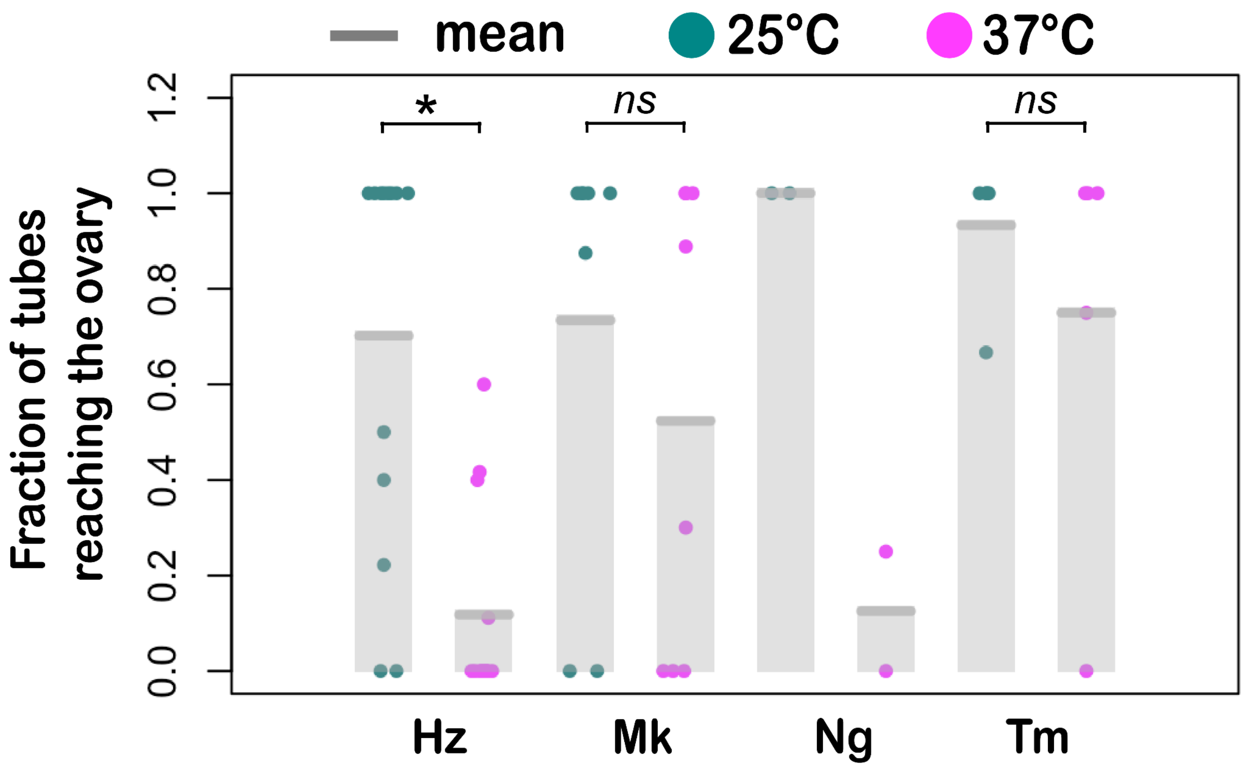


**Supplemental Figure S2. Fraction of tomato pollen tubes reaching the ovule at optimal and high temperatures under limited pollination conditions.** The fraction of pollen tubes, grown in the pistil under conditions highlighted in Figure 2C, that reached the ovules per genotype.

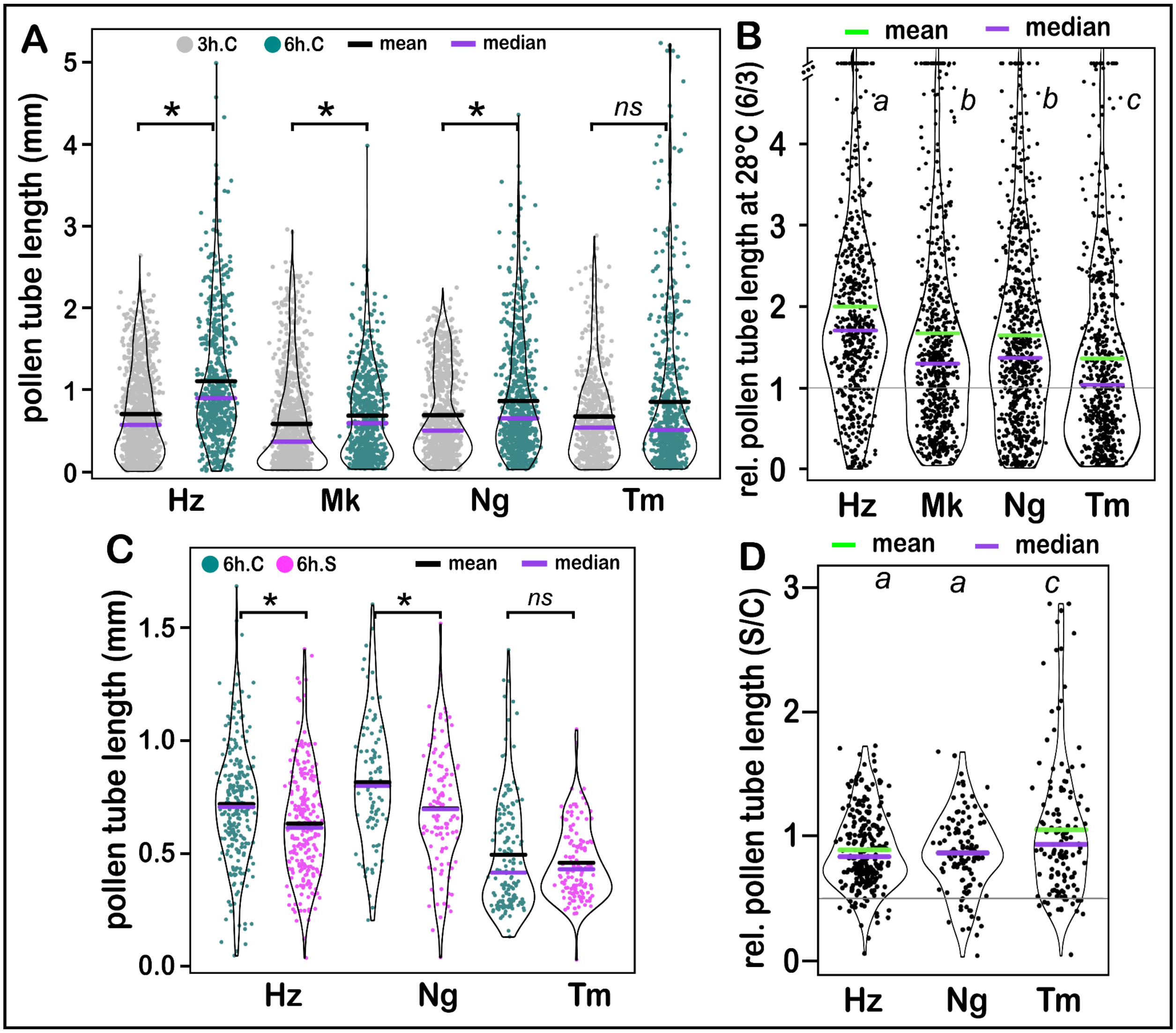


**Supplemental Figure S3. Effects of growth time and temperature on pollen tube elongation.** (A) pollen tube length after 3 hours (gray) and 6 hours (dark green) of growth at 25°C. (B) relative pollen tube length (3 hours/6 hours at 28°C). (C) repeat of figure 3 using pollen tubes grown on petri dishes instead of conical test tubes as done in figure 3 and for the RNA seq. (D) Relative high temperature pollen tube length of pollen from different varieties.

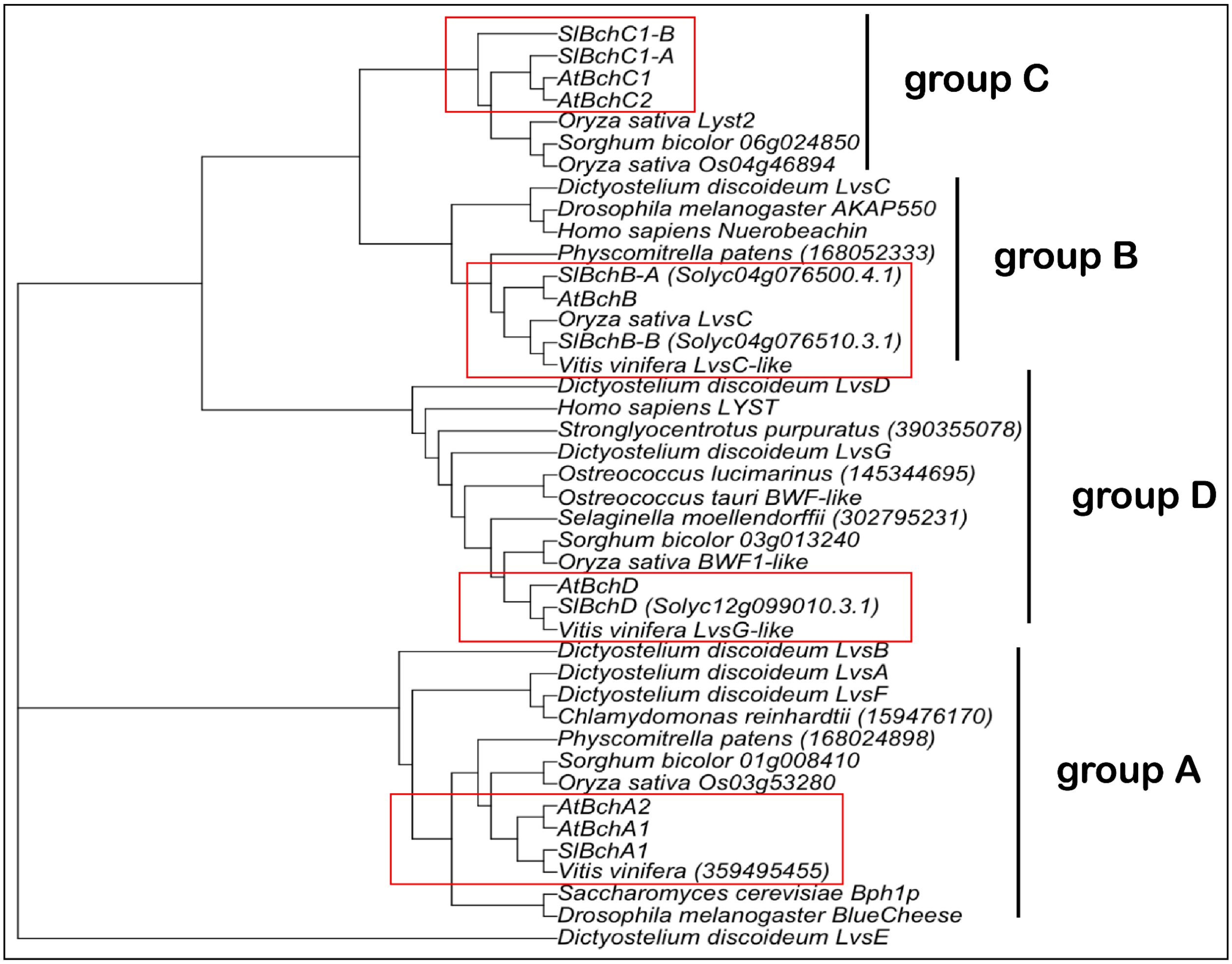


**Supplemental Figure S4. Phylogenetic tree of tomato Spirrig Homologs.** We used the tomato genes annotated as beach domain-containing homologs and the phylogenetic tree previously reported by Teh et al. [1] to finetune the annotation of tomato Spirrig homologs. The tree was constructed using a rooted neighbor joining method.

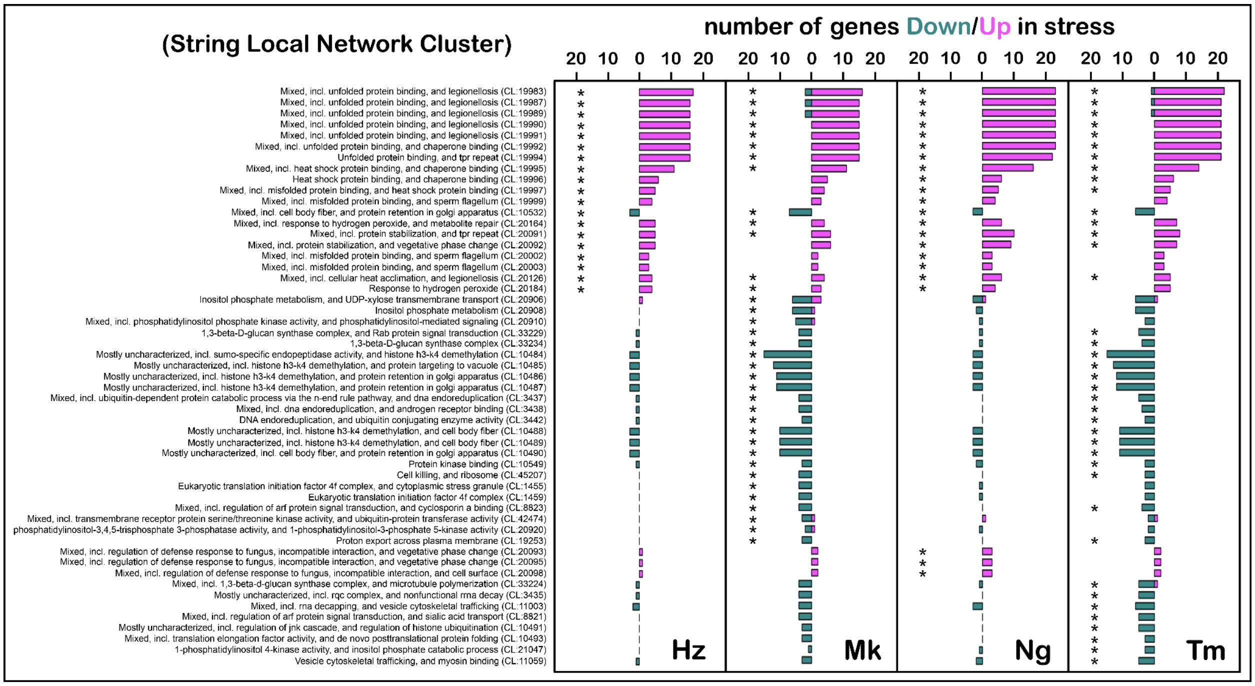


**Supplemental Figure S5. Gene set enrichment analysis (String local networks).** String local networks that are enriched in at least one cultivar due to heat stress during in-vitro pollen tube growth.

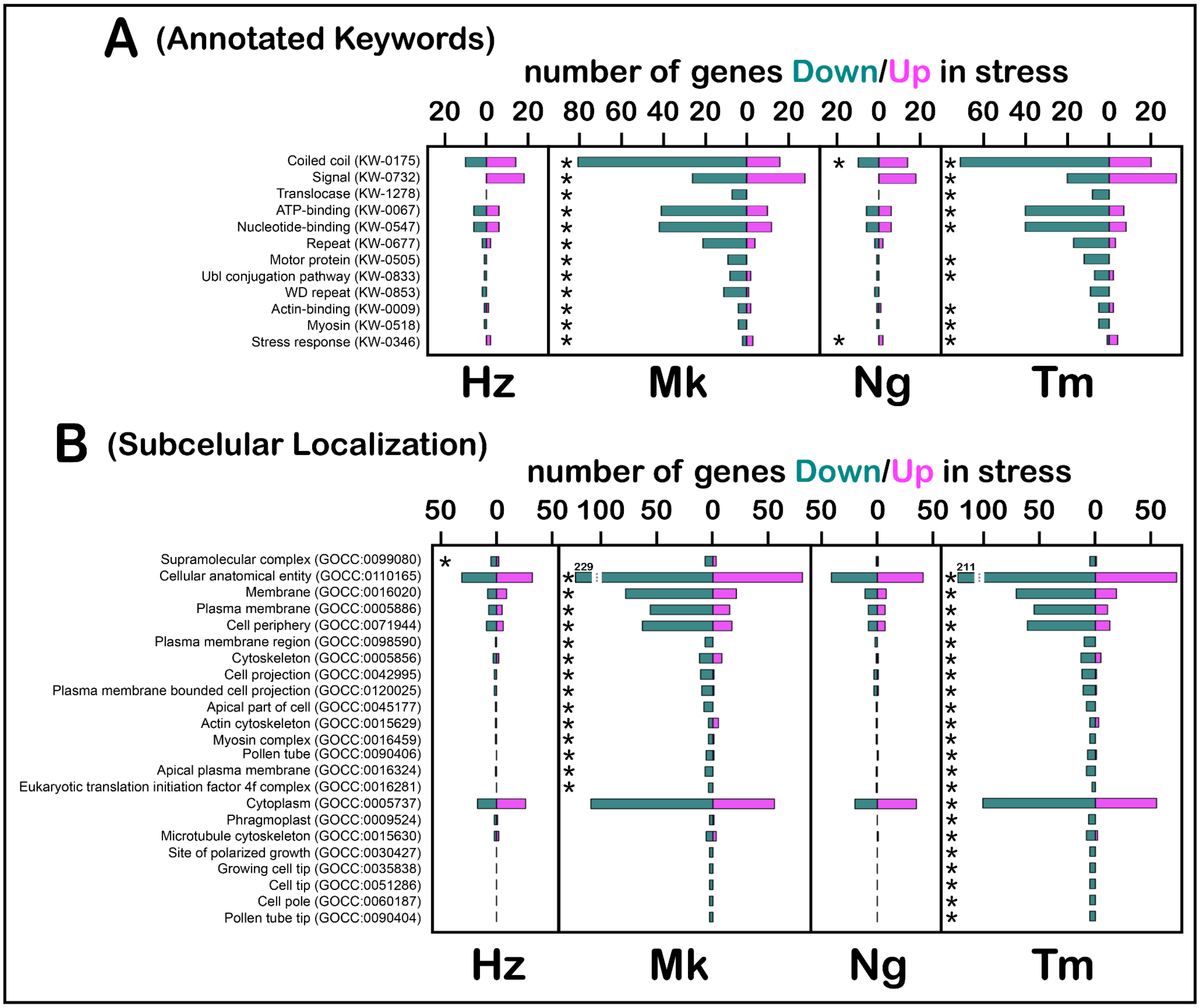


**Supplemental Figure S6. Gene set enrichment analysis (annotated keywords, subcellular localization).** Annotated keywords (A) and cellular compartments (B) enriched in at least one cultivar due to heat stress during in-vitro pollen tube growth.

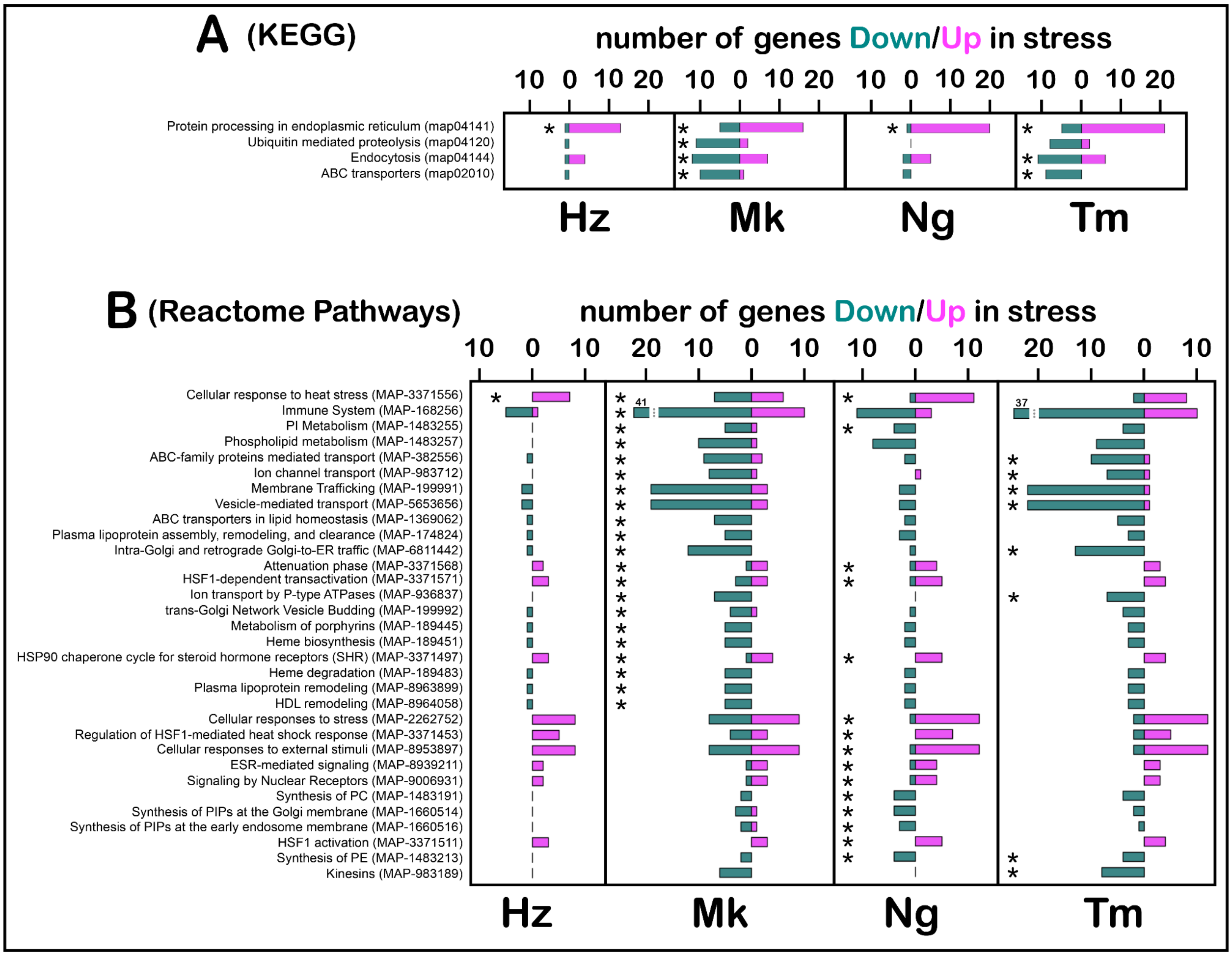


**Supplemental Figure S7. Gene set enrichment analysis (KEGG, Reactome).** Annotated (A) and Reactome (B) pathways enriched in at least one cultivar due to heat stress during in-vitro pollen tube growth.

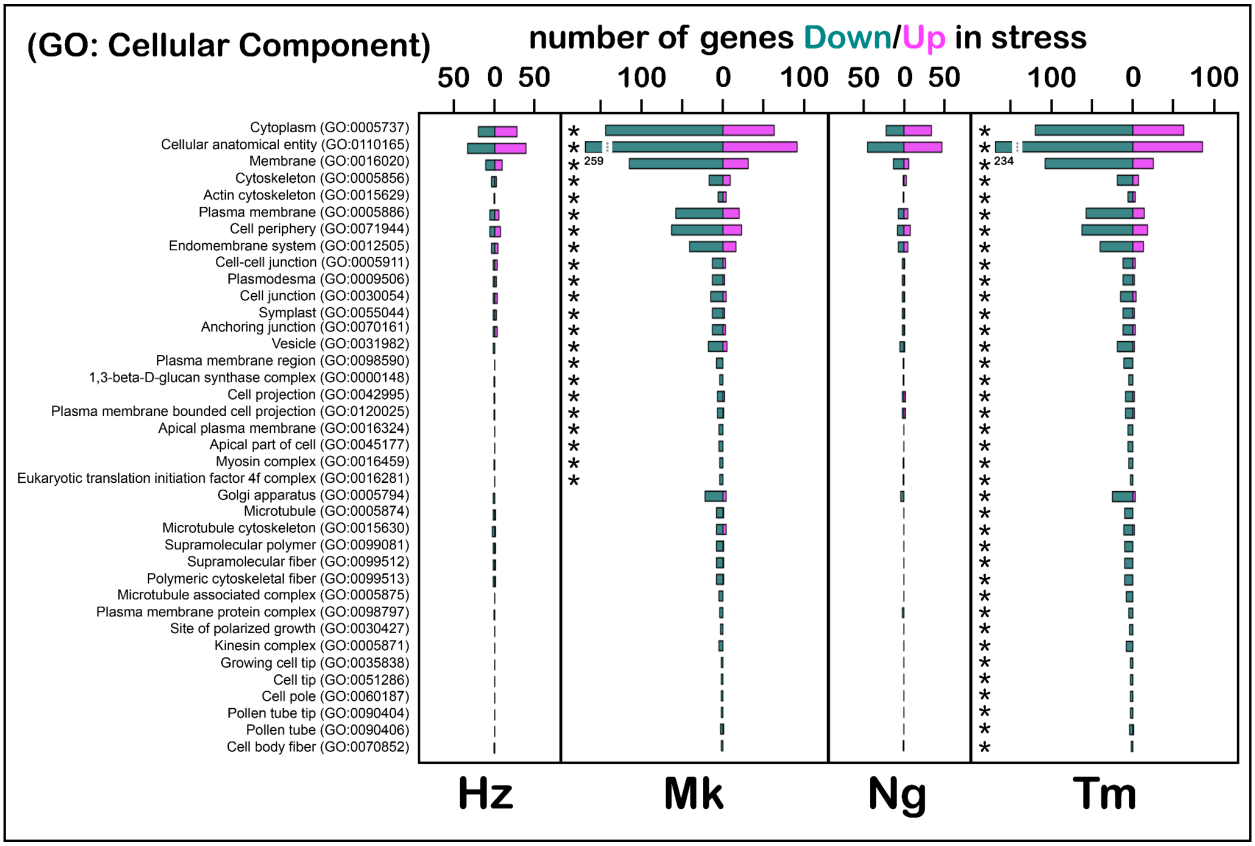


**Supplemental Figure S8. Gene set enrichment analysis (Cellular Component).** Cellular components enriched in at least one cultivar due to heat stress during in-vitro pollen tube growth.

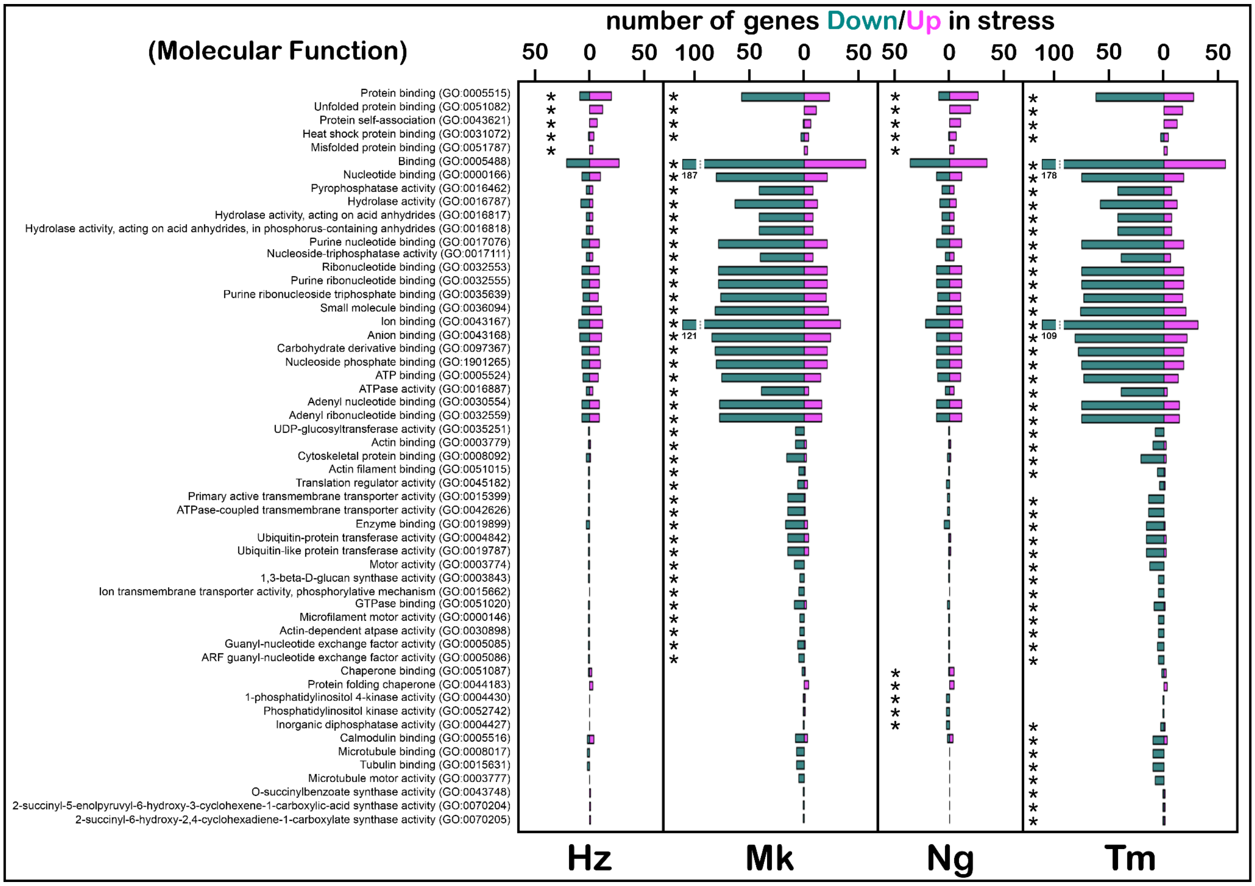


**Supplemental Figure S9. Gene set enrichment analysis (Molecular Function).** Molecular functions enriched in at least one cultivar due to heat stress during in-vitro pollen tube growth.

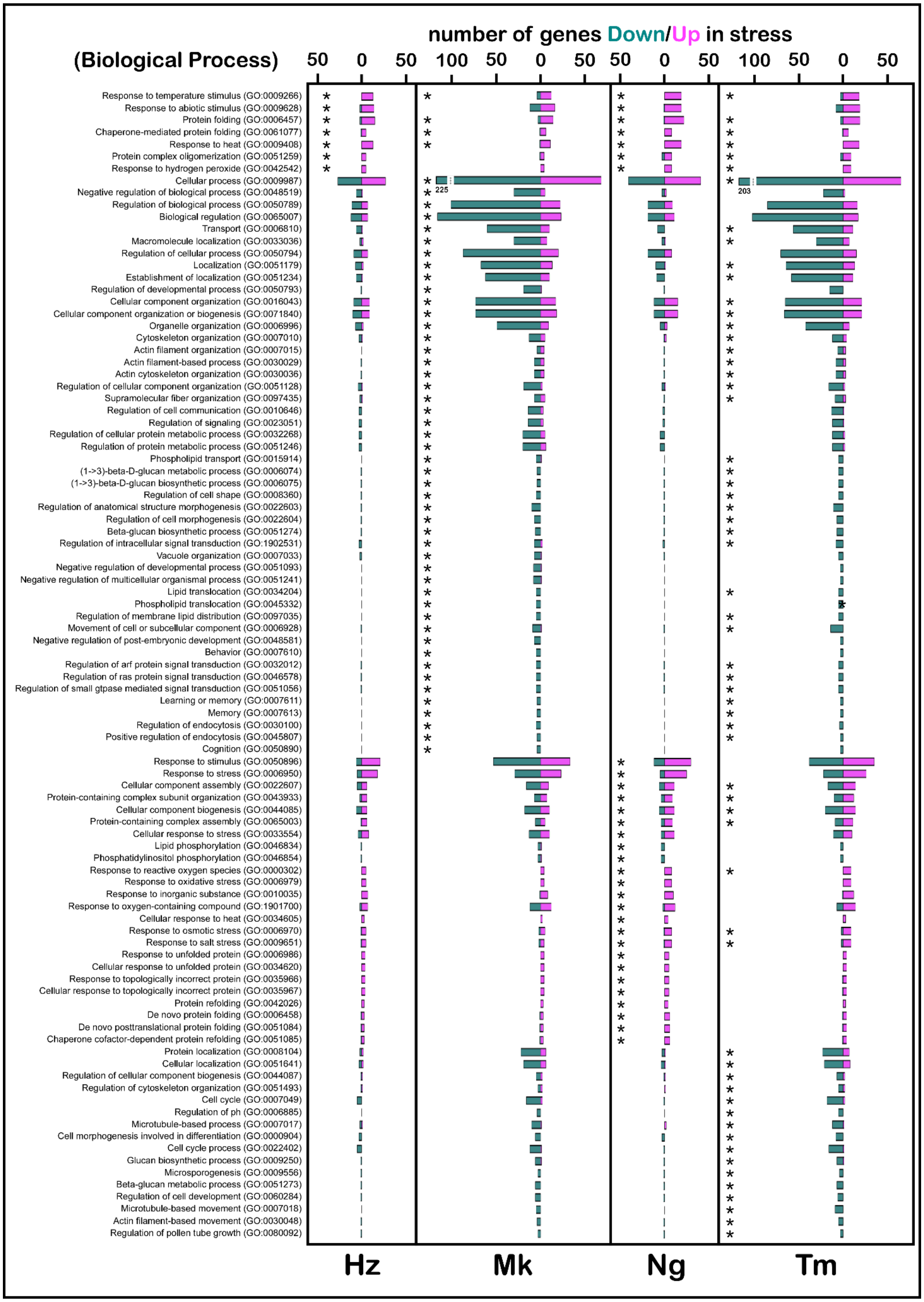


**Supplemental Figure S10. Gene set enrichment analysis (Biological Process).** Biological processes enriched in at least one cultivar due to heat stress during in-vitro pollen tube growth.

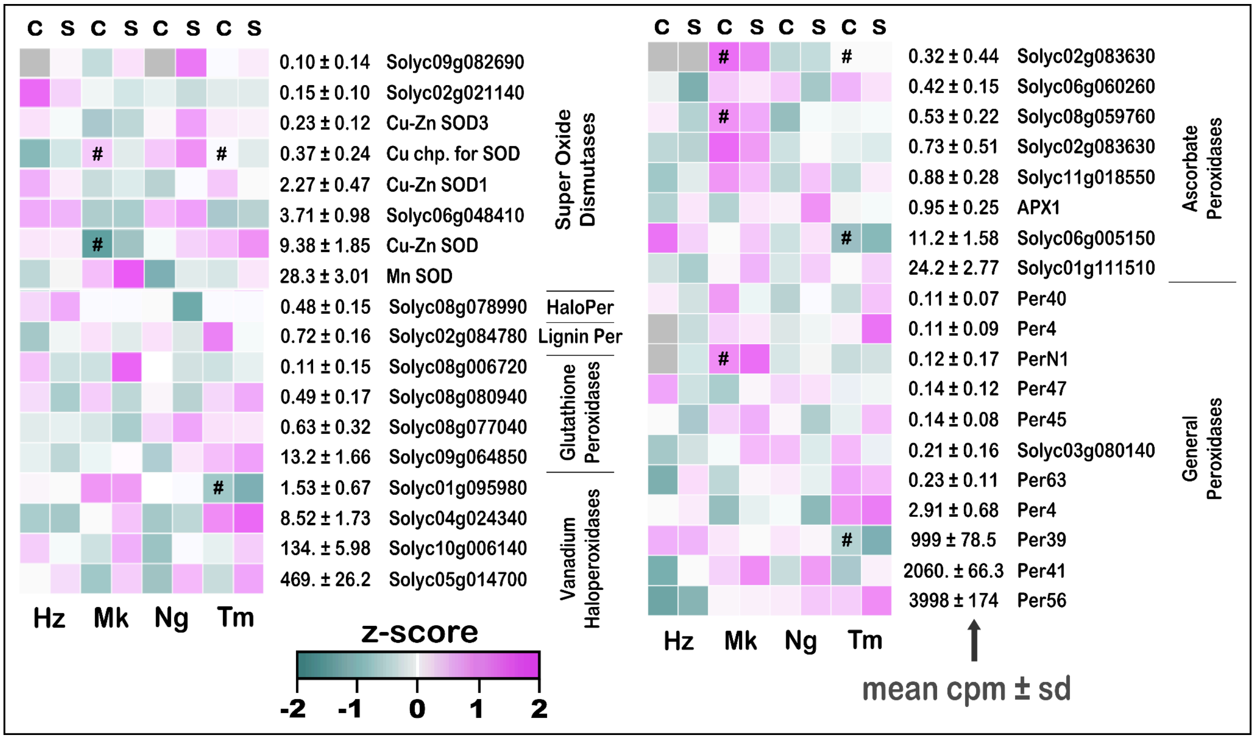


**Supplemental Figure S11. Heat maps superoxide dismutases and peroxidases genes expressed in growing pollen tubes.** * = significant compared to the same cultivar's 28°C treatment; # = significant compared to Heinz at 28°C. Heatmaps display Z-scores, the number of standard deviations above (magenta), below (dark green), or at the mean (white) CPM across all treatments for each gene. Hz, Heinz; Mk, Malintka; Ng, Nagcarlang; Tm, Tamaulipas; S, Stress (28°C, 3 hours; 37°C, 3 hours ); C, Control (28°C, 6 hours).

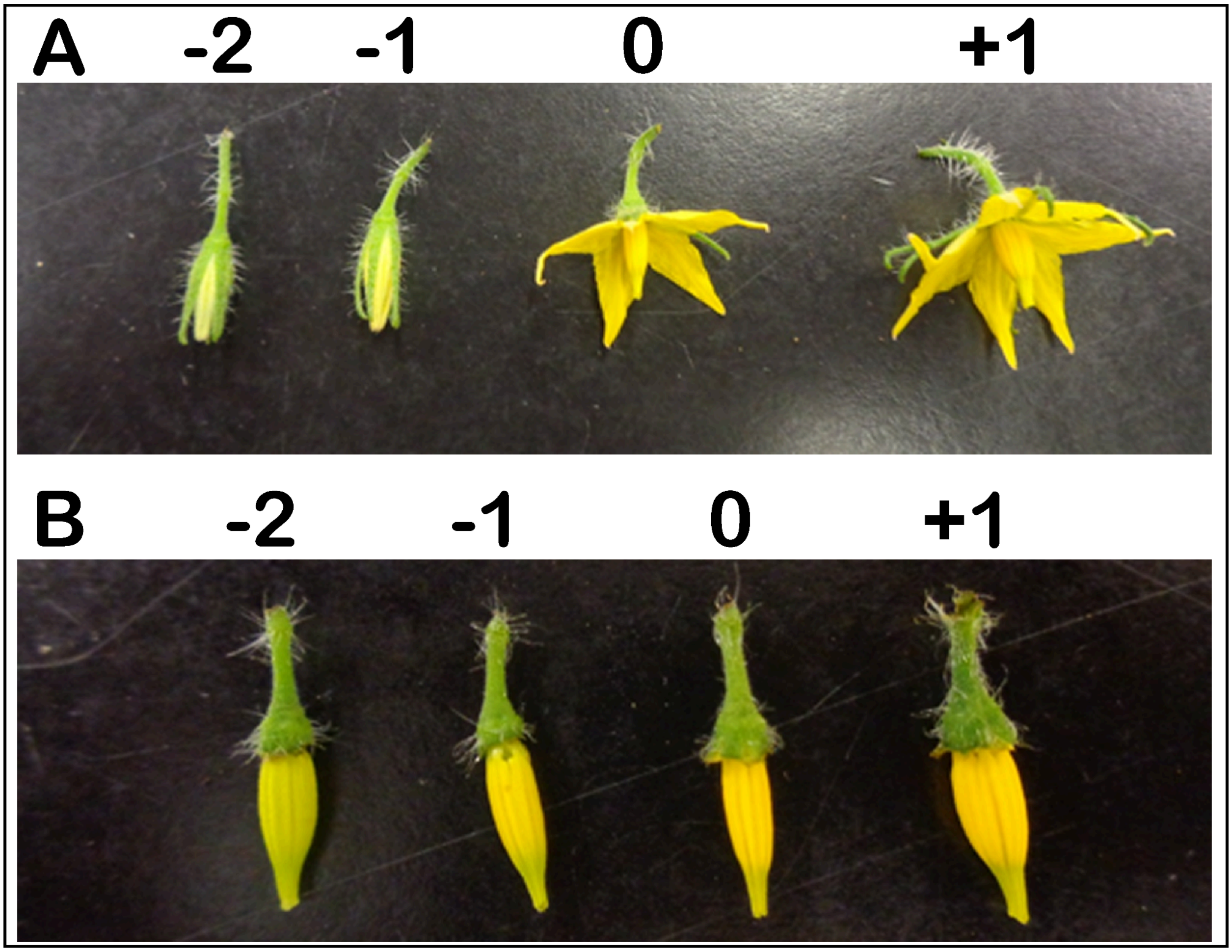


**Supplemental Figure S12. Tomato flower staging.** Tomato flower (A) and anther cone (B) appearances at different flower development stages. Heinz flowers are shown.

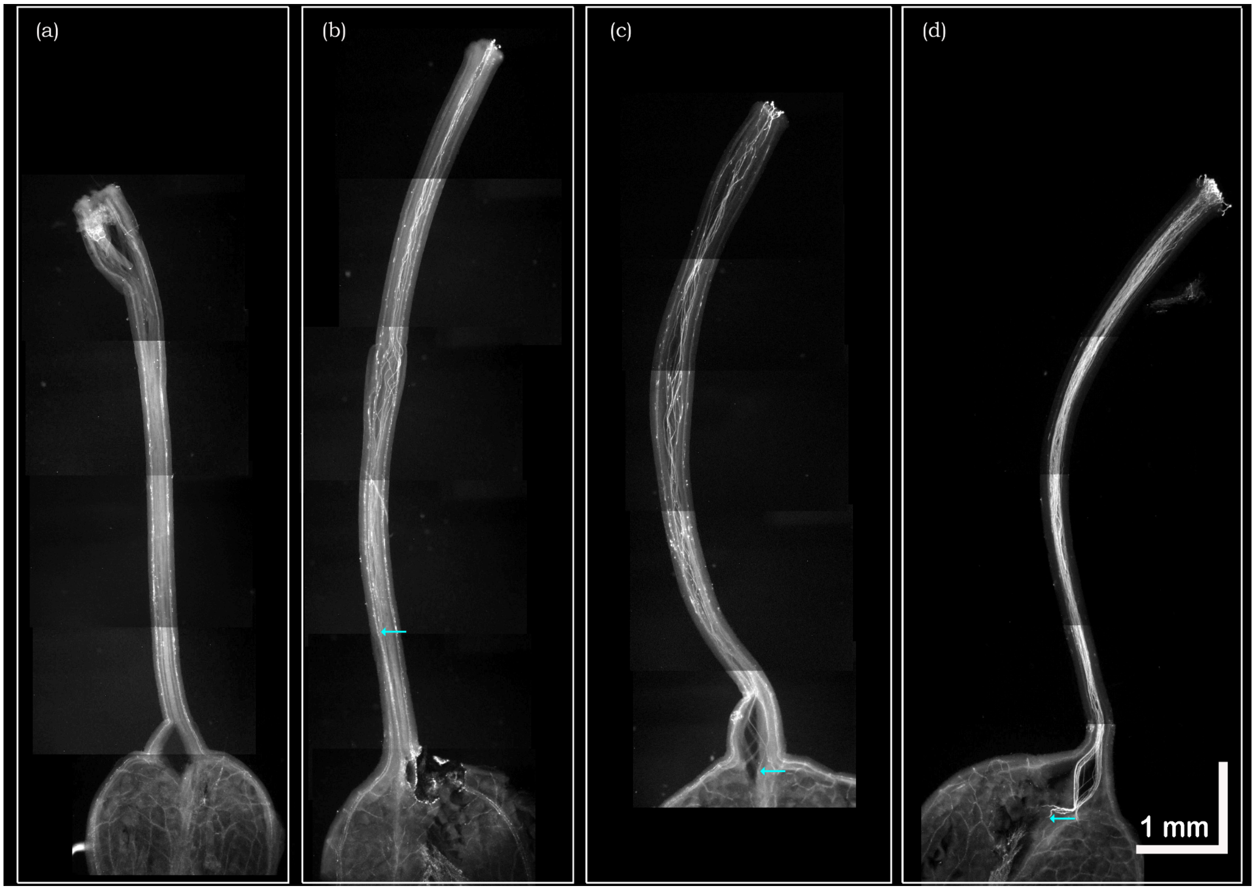


**Supplemental Figure S13. Analysis of pollen tube growth in the pistil. (A-D)** Examples of aniline blue-stained pistils with limited pollinations. The position of the tip of the longest pollen tube measured is indicated with an arrow.

Supplemental Table S1. Cultivars noted for thermtolerant fruit production

| TGRC Accession # | Name | Comments on cultivars from UC Davis C.M. Rick Tomato Genetics Resource Center (https://tgrc.ucdavis.edu/) | Refs. |
|---|---|---|---|
| | | **Reference Cultivar** | |
| LA4345 | *Heinz | This line was used for the tomato genome sequencing project. See variety description in TGC 54: 26. Given name: Heinz 1706 - BG | 1 |
| | | **Cultivars noted for thermotolerant fruit set** | |
| LA1994 | *Tamaulipas | *L. esculentum* primitive cultivar with small yellow fruits. Noted to have “..production abilities under high temperatures”. Named in this study after the location where it was first identified. | 2,3 |
| LA3120 | *Malintka | Heat and stress tolerant. Given name: Malintka 101 | 4,5 |
| LA2661 | *Nagcarlang | Heat and stress tolerant | 6 |
| LA2661 | Gold Nugget | Fruit are parthenocarpic under low (and possibly high) temperatures (ref. Baggett and Kean 1985 HortSci. 20: 957). Yellow fruit color in this line is assumed to be controlled by the *r* gene; production of lycopene is inhibited | 7 |
| LA3320 | Hotset | Heat tolerance | |
| LA2375 | San Marzano | Originally listed as "Lc", but does not contain the Lc gene. From USDA-GRIN: "It is a special selection of San Marzano, carrying strong resistance to heat sterility." | |
| LA2662 | Saladette | Heat tolerance; bred by Paul Leeper (TGC 27). "Small sp, c [our stock is C], I, S. Fruit 40-60g, plum to globe shaped, firm 2-3 locules... Sets under high temperatures / high humidity." | |

* cultivars used in this study

| Supplementary Table S18. Accessions used to construct phylogeny presented in Figure 1A. | | | |
|---|---|---|---|
| **SRAID** | **Name** | **LA Accession #** | **Original Study** |
| SRR1572498 | TS-95 Moneymaker | | [1] |
| SRR11205483 | BGV006865 Nanopore 2 | | [2] |
| SRR11205474 | Brandywine Illumina | | [2] |
| SRR11205476 | Floradade Illumina | LA3242 | [2] |
| SRR11195634 | PAS014479 Illumina | | [2] |
| SRR1572398 | TS-230 Cerise Ildi 1 | | [1] |
| SRR1572652 | TS-278 Early Santa Clara 1 | LA0517 | [1] |
| SRR1572473 | TS-49 Earliana 2 | LA3238 | [1] |
| SRR1572613 | TS-228# M-82 | LA3475 | [1] |
| SRR1572461 | TS-10 Marglobe | LA0502 | [1] |
| SRR1572358 | TS-98 Gold Nugget 1 | LA4355 | [1] |
| SRR1572432 | TS-295 Puna 2 | LA0417 | [1] |
| SRR1572591 | TS-200 Hot set 2 | LA3320 | [1] |
| SRR1572480 | TS-68 Chiclayo 1 | LA0395 | [1] |
| SRR1572384 | Nagcarlang.Genome | LA2661 | [1] |
| SRR1572375 | Malintka.Genome | LA3120 | [1] |
| SRR1572593 | TS-201 San Salvador | LA1210 | [1] |
| SRR1572452 | TS-1 VF-36 | LA0490 | [1] |
| SRR1572460 | TS-9 Ailsa Craig | LA2838A | [1] |
| SRR1572457 | TS-6 San Marzano | LA3008 | [1] |
| SRX25048037 | Tamaulipas WGS | LA1994 | This Study |
| SRR1572579 | TS-188 Stone 1 | LA1506 | [1] |
| SRR1572377 | TS-149 Villa Hermosa | LA1425 | [1] |
| SRR1572419 | TS-280 Cerese du sud ouest N 2 | | [1] |
| SRR1572322 | TS-39 Cerise Gold 1 | | [1] |
| SRR1572438 | TS-301 Santa Cruz near Shintuyo | LA2688 | [1] |
| SRR1572440 | TS-303 Los Banos 1 | LA1461 | [1] |
| SRR1572378 | TS-154 Muna | LA1623 | [1] |
| SRR1572380 | TS-165 Veracruz | | [1] |
| SRR1572558 | TS-167 Tegucigalpa | LA0147 | [1] |
| SRR1572541 | TS-150 Tarapoto 2 | | [1] |
| SRR1572300 | TS-27 N135 Green Gage 1 | | [1] |
| SRR1572325 | TS-40 Cerise VFNT 2 | | [1] |
| SRR1572303 | TS-28 N 347 Yablochnyi 2 | | [1] |
| SRR1572411 | TS-257 Marpha N2 1 | | [1] |
| SRR1572345 | TS-75 Poire jaune | | [1] |

| | | | |
|---|---|---|---|
| SRR1572404 | TS-240 Arroyo Rico | | [1] |
| SRR1572459 | TS-8 E-6203 | | [1] |

| **Supplement Table S19: P-values obtained for phenotypic analyses.** | | | |
|---|---|---|---|
| groups | test | p.value | figure |
| Hz_C, Hz_S | Mann-Whitney test | 0.0001681995159 | 1C |
| Ng_C, Ng_S | Mann-Whitney test | 8.53E-05 | 1C |
| Tm_C, Tm_S | Mann-Whitney test | 0.1067235999 | 1C |
| Hz_S, Ng_S, Tm_S | Kruskal-Wallis test | 0.04487087313 | 1D |
| Hz_S, Ng_S | Dunn test | 0.2054926 | 1D |
| Hz_S, Tm_S | Dunn test | 0.02818885 | 1D |
| Ng_S, Tm_S | Dunn test | 0.01060241 | 1D |
| Hz_C, Hz_S | Mann-Whitney test | 0.01225221936 | 1E |
| Ng_C, Ng_S | Mann-Whitney test | 0.02518943906 | 1E |
| Tm_C, Tm_S | Mann-Whitney test | 0.02074709026 | 1E |
| Hz_S, Ng_S, Tm_S | Kruskal-Wallis test | 0.3130893415 | 1F |
| Hz_S, Ng_S | Dunn test | 0.06965967 | 1F |
| Hz_S, Tm_S | Dunn test | 0.1233893 | 1F |
| Ng_S, Tm_S | Dunn test | 0.4335576 | 1F |
| Hz_C, Hz_S | Mann-Whitney test | 0.03381771792 | 2B |
| Ng_C, Ng_S | Mann-Whitney test | 0.002778685068 | 2B |
| Tm_C, Tm_S | Mann-Whitney test | 0.1693272972 | 2B |
| Hz_C, Hz_S | Mann-Whitney test | 1.21E-17 | 2D |
| Mk_C, Mk_S | Mann-Whitney test | 0.001178195502 | 2D |
| Ng_C, Ng_S | Mann-Whitney test | 0.0006167260896 | 2D |
| Tm_C, Tm_S | Mann-Whitney test | 0.002564108215 | 2D |
| Hz_S, Mk_S, Ng_S, Tm_S | Kruskal-Wallis test | 3.23E-10 | 2E |
| Hz_S, Mk_S | Dunn test | 3.01E-06 | 2E |
| Hz_S, Ng_S | Dunn test | 0.002573269 | 2E |
| Hz_S, Tm_S | Dunn test | 4.78E-09 | 2E |
| Mk_S, Ng_S | Dunn test | 0.3443322 | 2E |
| Mk_S, Tm_S | Dunn test | 0.01471903 | 2E |
| Ng_S, Tm_S | Dunn test | 0.001208162 | 2E |
| Hz.C, Hz.S | Mann-Whitney test | 3.98E-38 | 3C |
| Mk.C, Mk.S | Mann-Whitney test | 0.001049014664 | 3C |

| | | | |
|---|---|---|---|
| Ng.C, Ng.S | Mann-Whitney test | 8.83E-13 | 3C |
| Tm.C, Tm.S | Mann-Whitney test | 0.121559918 | 3C |
| Hz.S, Mk.S, Ng.S, Tm.S | Kruskal-Wallis test | 3.02E-30 | 3D |
| Hz.S, Mk.S | Dunn test | 6.75E-21 | 3D |
| Hz.S, Ng.S | Dunn test | 2.87E-11 | 3D |
| Hz.S, Tm.S | Dunn test | 3.29E-26 | 3D |
| Mk.S, Ng.S | Dunn test | 0.0008688196 | 3D |
| Mk.S, Tm.S | Dunn test | 0.08734591 | 3D |
| Ng.S, Tm.S | Dunn test | 3.40E-07 | 3D |
| lk_C, Mk_S, Ng_C, Ng_S | Kruskal-Wallis test | 1.25E-09 | 7B |
| Hz_C, Hz_S | Dunn test | 0.0005908362 | 7B |
| Hz_C, Mk_C | Dunn test | 0.3500682 | 7B |
| Hz_C, Mk_S | Dunn test | 0.0005848402 | 7B |
| Hz_C, Ng_C | Dunn test | 0.02158997 | 7B |
| Hz_C, Ng_S | Dunn test | 0.0001883232 | 7B |
| Hz_C, Tm_C | Dunn test | 0.03699531 | 7B |
| Hz_C, Tm_S | Dunn test | 0.0001883232 | 7B |
| Hz_S, Mk_C | Dunn test | 0.000427551 | 7B |
| Hz_S, Mk_S | Dunn test | 0.02370062 | 7B |
| Hz_S, Ng_C | Dunn test | 0.000427551 | 7B |
| Hz_S, Ng_S | Dunn test | 0.384849 | 7B |
| Hz_S, Tm_C | Dunn test | 0.0005968727 | 7B |
| Hz_S, Tm_S | Dunn test | 0.02021199 | 7B |
| Mk_C, Mk_S | Dunn test | 0.0004237934 | 7B |
| Mk_C, Ng_C | Dunn test | 0.0189886 | 7B |
| Mk_C, Ng_S | Dunn test | 0.0001637069 | 7B |
| Mk_C, Tm_C | Dunn test | 0.03375408 | 7B |
| Mk_C, Tm_S | Dunn test | 0.0001637069 | 7B |
| Mk_S, Ng_C | Dunn test | 0.0004237934 | 7B |
| Mk_S, Ng_S | Dunn test | 0.05909716 | 7B |
| Mk_S, Tm_C | Dunn test | 0.0005908362 | 7B |

| | | | |
|---|---|---|---|
| Mk_S, Tm_S | Dunn test | 0.0003157111 | 7B |
| Ng_C, Ng_S | Dunn test | 0.0001192817 | 7B |
| Ng_C, Tm_C | Dunn test | 0.2818514 | 7B |
| Ng_C, Tm_S | Dunn test | 0.0001192817 | 7B |
| Ng_S, Tm_C | Dunn test | 0.0001896423 | 7B |
| Ng_S, Tm_S | Dunn test | 0.0246831 | 7B |
| Tm_C, Tm_S | Dunn test | 0.0001896423 | 7B |
| Mk.C, Mk.S, Ng.C, Ng.S, [illegible] | Kruskal-Wallis test | 1.49E-95 | 8C |
| Hz.C, Hz.S | Dunn test | 4.18E-14 | 8C |
| Hz.C, Mk.C | Dunn test | 0.1161289 | 8C |
| Hz.C, Mk.S | Dunn test | 6.67E-12 | 8C |
| Hz.C, Ng.C | Dunn test | 3.72E-17 | 8C |
| Hz.C, Ng.S | Dunn test | 6.09E-33 | 8C |
| Hz.C, Tm.C | Dunn test | 5.01E-33 | 8C |
| Hz.C, Tm.S | Dunn test | 1.77E-11 | 8C |
| Hz.S, Mk.C | Dunn test | 0.002057203 | 8C |
| Hz.S, Mk.S | Dunn test | 0.0005726218 | 8C |
| Hz.S, Ng.C | Dunn test | 0.001304948 | 8C |
| Hz.S, Ng.S | Dunn test | 2.19E-13 | 8C |
| Hz.S, Tm.C | Dunn test | 4.14E-46 | 8C |
| Hz.S, Tm.S | Dunn test | 1.13E-25 | 8C |
| Mk.C, Mk.S | Dunn test | 4.61E-05 | 8C |
| Mk.C, Ng.C | Dunn test | 8.58E-06 | 8C |
| Mk.C, Ng.S | Dunn test | 1.01E-11 | 8C |
| Mk.C, Tm.C | Dunn test | 1.13E-19 | 8C |
| Mk.C, Tm.S | Dunn test | 1.46E-07 | 8C |
| Mk.S, Ng.C | Dunn test | 0.2035328 | 8C |
| Mk.S, Ng.S | Dunn test | 0.05147226 | 8C |
| Mk.S, Tm.C | Dunn test | 1.14E-26 | 8C |
| Mk.S, Tm.S | Dunn test | 5.20E-17 | 8C |
| Ng.C, Ng.S | Dunn test | 4.52E-05 | 8C |

| | | | |
|---|---|---|---|
| Ng.C, Tm.C | Dunn test | 4.43E-33 | 8C |
| Ng.C, Tm.S | Dunn test | 1.53E-25 | 8C |
| Ng.S, Tm.C | Dunn test | 3.20E-40 | 8C |
| Ng.S, Tm.S | Dunn test | 1.20E-34 | 8C |
| Tm.C, Tm.S | Dunn test | 1.70E-08 | 8C |
| Hz.S, Mk.S, Ng.S, Tm.S | Kruskal-Wallis test | 0.003263110118 | 8D |
| Hz.S, Mk.S | Dunn test | 0.05291903 | 8D |
| Hz.S, Ng.S | Dunn test | 0.004830631 | 8D |
| Hz.S, Tm.S | Dunn test | 0.1372178 | 8D |
| Mk.S, Ng.S | Dunn test | 0.0001984515 | 8D |
| Mk.S, Tm.S | Dunn test | 0.3039269 | 8D |
| Ng.S, Tm.S | Dunn test | 0.001277665 | 8D |
| Mk.C, Mk.S, Ng.C, Ng.S, | Kruskal-Wallis test | 4.20E-17 | 8E |
| Hz.C, Hz.S | Dunn test | 0.001468677 | 8E |
| Hz.C, Mk.C | Dunn test | 0.004600559 | 8E |
| Hz.C, Mk.S | Dunn test | 2.90E-08 | 8E |
| Hz.C, Ng.C | Dunn test | 0.2151515 | 8E |
| Hz.C, Ng.S | Dunn test | 0.0005441994 | 8E |
| Hz.C, Tm.C | Dunn test | 4.21E-12 | 8E |
| Hz.C, Tm.S | Dunn test | 6.09E-11 | 8E |
| Hz.S, Mk.C | Dunn test | 0.1777218 | 8E |
| Hz.S, Mk.S | Dunn test | 0.0001347083 | 8E |
| Hz.S, Ng.C | Dunn test | 0.06966104 | 8E |
| Hz.S, Ng.S | Dunn test | 0.1982998 | 8E |
| Hz.S, Tm.C | Dunn test | 4.74E-07 | 8E |
| Hz.S, Tm.S | Dunn test | 1.80E-06 | 8E |
| Mk.C, Mk.S | Dunn test | 0.01749388 | 8E |
| Mk.C, Ng.C | Dunn test | 0.04350889 | 8E |
| Mk.C, Ng.S | Dunn test | 0.3354187 | 8E |
| Mk.C, Tm.C | Dunn test | 0.003480694 | 8E |
| Mk.C, Tm.S | Dunn test | 0.00278952 | 8E |

| | | | |
|---|---|---|---|
| Mk.S, Ng.C | Dunn test | 1.04E-05 | 8E |
| Mk.S, Ng.S | Dunn test | 0.002090235 | 8E |
| Mk.S, Tm.C | Dunn test | 0.3185331 | 8E |
| Mk.S, Tm.S | Dunn test | 0.272022 | 8E |
| Ng.C, Ng.S | Dunn test | 0.006046425 | 8E |
| Ng.C, Tm.C | Dunn test | 2.20E-08 | 8E |
| Ng.C, Tm.S | Dunn test | 1.62E-07 | 8E |
| Ng.S, Tm.C | Dunn test | 5.20E-05 | 8E |
| Ng.S, Tm.S | Dunn test | 0.000119682 | 8E |
| Tm.C, Tm.S | Dunn test | 0.4531784 | 8E |
| Hz.S, Mk.S, Ng.S, Tm.S | Kruskal-Wallis test | 0.01031398668 | 8F |
| Hz.S, Mk.S | Dunn test | 0.001642197 | 8F |
| Hz.S, Ng.S | Dunn test | 0.4666964 | 8F |
| Hz.S, Tm.S | Dunn test | 0.05702897 | 8F |
| Mk.S, Ng.S | Dunn test | 0.002373373 | 8F |
| Mk.S, Tm.S | Dunn test | 0.2241099 | 8F |
| Ng.S, Tm.S | Dunn test | 0.0255828 | 8F |
| Hz.C, Hz.S | Welch Two Sample t-test | 0.1649557582 | 8G |
| Mk.C, Mk.S | Welch Two Sample t-test | 0.1410985543 | 8G |
| Ng.C, Ng.S | Welch Two Sample t-test | 0.1277806314 | 8G |
| Tm.C, Tm.S | Welch Two Sample t-test | 0.7463963428 | 8G |
